# Genetic landscape of an *in vivo* protein interactome

**DOI:** 10.1101/2023.12.14.571726

**Authors:** Savandara Besse, Tatsuya Sakaguchi, Louis Gauthier, Zahra Sahaf, Olivier Péloquin, Lidice Gonzalez, Xavier Castellanos-Girouard, Nazli Koçatug, Chloé Matta, Julie G. Hussin, Stephen W. Michnick, Adrian W.R. Serohijos

## Abstract

Protein-protein interaction (PPI) networks accurately map environmental perturbations to their molecular consequences in cells, but effects of genome-wide genetic variation on PPIs remain unknown. We hypothesized that PPI networks integrate genetic and environmental effects, potentially defining biochemical mechanisms underlying complex polygenic traits. Here, we measured 61 PPIs in inbred strains of *Saccharomyces cerevisiae* with ∼12,000 single-nucleotide polymorphisms (SNPs) across the genome. Unlike mRNA expression and protein abundance that are primarily affected by SNPs local (in “cis”) to a gene, PPIs are predominantly affected by SNPs far (in “trans”) to the genomic loci of the interacting proteins. However, consistent with the PPI network’s small-world characteristic, these transacting SNPs are in neighboring genes in the network. We likewise discovered SNPs in non-coding RNAs and post-transcriptional regulators (3’ UTRs) with, counterintuitively, larger PPI-modulating effects than SNPs within protein-coding regions. Finally, we inferred known and novel mechanisms of action for yeast and human drugs.

**HIGHLIGHTS:**

- Protein-interaction quantitative trait locus (“piQTL”) mapping reveals sensitivity of *in vivo* PPIs to polymorphisms across the yeast genome
- Trans-piQTLs significantly outnumber and are stronger than cis-piQTLs
- SNPs in non-coding RNAs and 3’ UTRs have comparable effects to PPI as SNPs in coding regions
- piQTL mapping reveals known and novel mechanism of yeast and human drugs

## INTRODUCTION

A central question in biology is how genomic sequences influence complex traits and phenotypes. Genome-wide association studies (GWAS) suggest that genomic loci with significant statistical associations to the variation of complex or quantitative phenotypes (quantitative trait loci or QTLs) are spread across the genome. These include many genes without an obvious connection to the phenotype.^1-3^ There are now over 400,000 unique associations spanning over 5,000 traits in humans.^4,5^ However, despite advances, assigning function and causality to the resulting QTLs remains a challenge.^6^ Following the Central Dogma that biological information flows from DNA to RNA to Proteins,^7^ led to the development of various approaches to map genetic variation to intermediate molecular phenotypes. Gene expression has been used as a proxy molecular trait and a functional intermediate between genomic and phenotypic variation, thus identifying genomic loci that regulate the mRNA level of genes or “expression QTLs” (eQTLs).^8-10^ Several other types of association mapping strategies have been developed based on chromatin accessibility, DNA methylation, transcription factor binding, and metabolomics.

The ultimate consequences of the genotype-phenotype-environment relationship (G×E) are reflected in the steady-state levels, post-translational modifications, and subcellular localization of proteins and the other proteins that they interact with.^11-19^ Additionally, genotype-phenotype relationships based on protein abundances (“protein abundance QTL” or pQTL) better link genetic variation to molecular mechanisms underlying phenotypes than eQTL.^20^ eQTL, however, remains more broadly accessible than pQTL due to simpler material handling, lower cost and analytical simplicity of transcriptomics by RNAseq compared to proteomics by mass-spectrometry. Nevertheless, there is a consensus that the eventual ability to measure proteomic variation and relate it to genetics is a major goal in understanding the molecular mechanism of complex traits and polygenic phenotypes.^21^

In principle, protein-protein interactions (PPI) could be more accurate reporters of G×E than eQTLs or pQTLs since PPI variations capture the entire fate of the two genes that encode the two interacting proteins, including effects of their synthesis, degradation and post-translational modifications (Figure 1A),^12,15,18^ and thus, PPIs are mechanistically closest to phenotype. Several results support this hypothesis. First, *in vivo* PPI network is a dynamic quantification of the cell’s architecture. Studies that map the localization and interactions of human proteins, such as the OpenCell,^18^ have shown that spatial proximity not only recovers annotated biological function but can also predict of novel ones. Second, dynamic PPI formalizes the notion in biology that association, binding, and co-localization of molecules, including proteins, in the cell is the standard operational definition of “function”. Third, many studies showed that the dynamics of protein-protein interactions accurately map environmental perturbations, such as drug treatments^12^ or infections by viruses^22,23^, to their molecular consequences inside the cell. Fourth, PPIs are sensitive to genomic mutations. For example, 55 targeted mutations in a GTPase affects its interaction with 316 protein partners *in vivo*.^24^ Sahni et al. assayed thousands of protein-coding missense mutations associated to Mendelian human diseases and showed that two-thirds of them perturb PPIs.^25^

**Figure 1.**
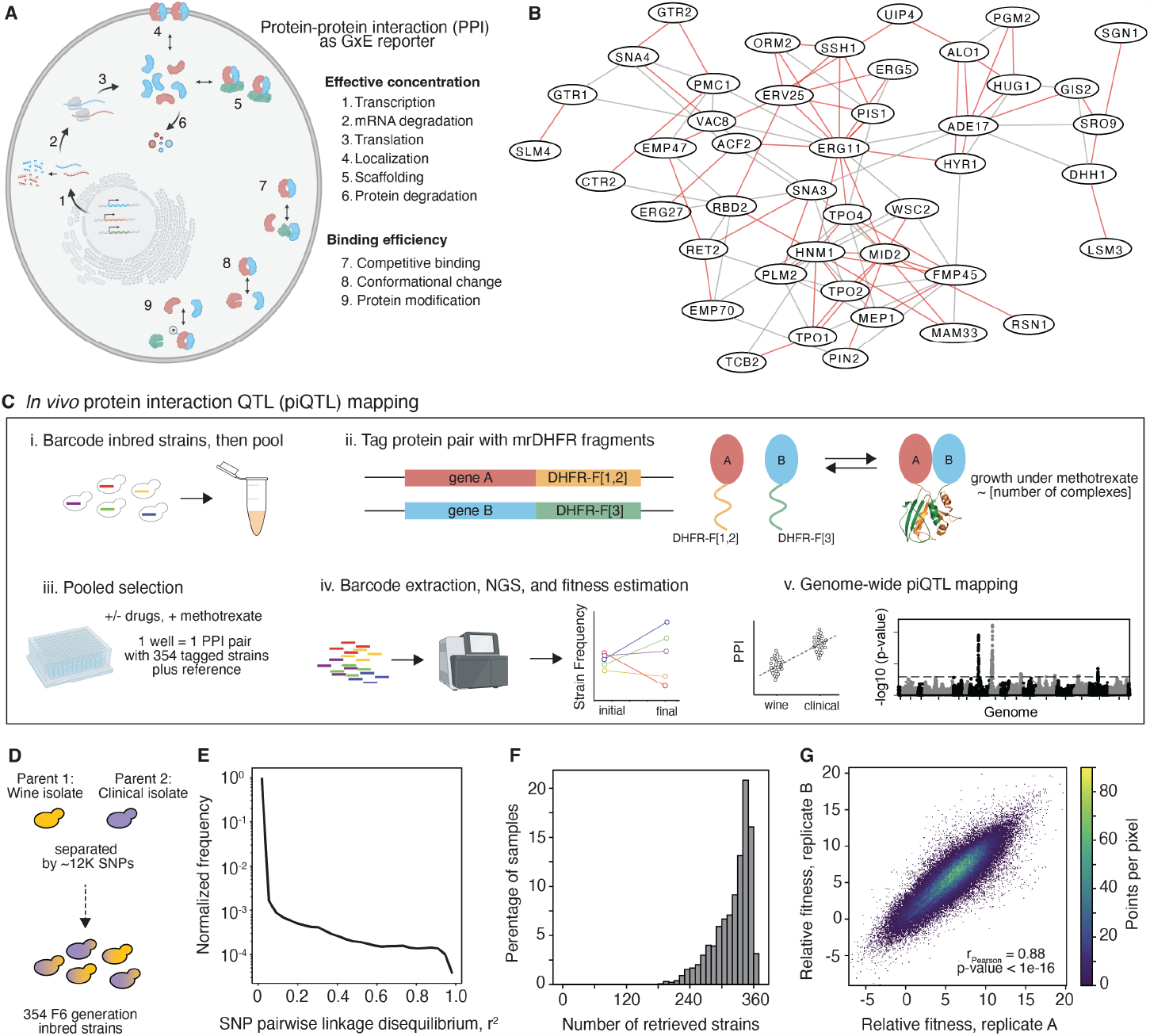
Protein-interaction QTL (piQTL) mapping in an inbred yeast strain cohort. **(A)** The strength of PPI is a consequence of many factors, some of which affect the concentration of binding pairs or their binding efficiency. **(B)** 61 PPIs spanning 44 unique genes chosen from diverse functional categories of biochemical pathways (Table S1) as reported in a previous PPI screen (all edges),^11^ and measured in this study (red edges). **(C)** Pooled screening strategy for *in vivo* piQTL mapping. *i*, Strains are individually barcoded in a neutral chromosomal locus by CRISPR/Cas9 and homologous recombination and then pooled (Figure S2A,B). *ii*, Step-wise editing on the pooled strains introduce the coding sequences for complementary fragments of methotrexate (MTX)-resistant (mr) mouse mrDHFR N-terminal (DHFR-F[1,2]) and C-terminal-fragment (DHFR-F[3]), 5’ to the ORFs, resulting in C-terminal fusions of the fragments to the two proteins of interest (Figure S2D,E). Reconstitution of mrDHFR due to interaction between the two proteins confers resistance against MTX. Strength of PPI is proportional to growth rate in MTX.^11,12^ *iii*, We propagated the pooled strains (354 inbred strains labeled with mrDHFR fragments and another 2 strains unlabeled to serve as reference for “no PPI”). Some strains were labeled with 2 distinct barcodes to determine within-well replicability (Figure S3C). Selection was performed in MTX as well as under drug perturbations. *iv*, Barcodes are amplified from the genome and quantified by NGS. Fitness of each strain and, consequently, the magnitude of PPIs, are estimated from changes in barcode frequencies during selection. *v*, Genome-wide piQTL mapping is performed using linear regression between PPI and genetic composition at each SNP locus. **(D** and **E)**, A collection of 354 yeast strains from F6 generation of inbreeding of wine and clinical isolates from a GWAS study.^26^ Inbreeding results in minimal pairwise linkage disequilibrium (LD) among ∼12,000 SNPs. **(F)** Number of strains recovered for all PPI and drug conditions in the pooled PPI screen indicate sufficient power for QTL mapping (Figure S1C). **(G)** Reproducibility of fitness estimates in biological replicates. See also Figures S1, S2, and S3.

Despite these works showing PPI to be an accurate reporter of environmental perturbations to cellular structure, the systematic effects of genome-wide genetic variation on PPI networks have never been measured. Here, we report how genetic variations correlate to changes in *in vivo* PPI using the budding yeast *Saccharomyces cerevisiae* (henceforth yeast) as a model system, and then subsequently, probe G×E relationships of drugs that target specific biochemical pathways in yeast and human diseases.

## RESULTS

### Pooled quantification of *in vivo* PPI in an inbred yeast cohort

We used a collection of 354 haploid yeast strains used in a previous GWAS study that resulted from the inbreeding of two divergent strains for 6 generations leading to a cohort with minimal linkage disequilibrium (LD) (Figure 1D,E; Figure S1A).^26^ The population contains ∼12,000 high-frequency SNPs spread across the 16 chromosomes and mitochondrial genome with at least 1 SNP per kilobase covering all gene elements including promoters, protein-coding, and other regulatory domains (Figure S1A, B). They are also found in non-coding RNAs and functional non-transcribed regions (Figure S1B). Based on power calculations (Figure S1C), our sample size of 354 inbred strains is well-above the statistical requirement to perform QTL studies, comparable to pioneering eQTL studies in mice (∼100 genomes)^27^ and Drosophila (∼200 genomes)^28,29^.

To measure the strength of PPI *in vivo*, we used a Protein-fragment Complementation Assay (PCA) based on the reporter enzyme methotrexate (MTX)-resistant murine dihydrofolate reductase (mrDHFR) (Figure 1C(ii)).^30^ mrDHFR PCA is ideal because it has been applied to determine the *in vivo* protein interactome of *S. cerevisiae*^11^ and to map environmental perturbation by drugs to known and novel biochemical pathways.^11-13,31,32^

In each of the 354 inbred strains, we integrated complementary N- and C-terminal fragments encoding for mrDHFR into the 3’-ends of the ORFs coding for the interacting protein pair. When the two tagged proteins interact, they bring the complementary mrDHFR fragments into proximity (∼8 nm)^30^ to fold into an active enzyme, which confers growth in cells in which the endogenous DHFR is inhibited by MTX.^11^ We measured the PPI of 61 protein pairs that cover 44 unique genes chosen based on diverse coverage of molecular function, biological processes, and cellular localization and which have been validated in a proteome-wide mrDHFR PCA screen^11^ (Figure 1B and Table S1(GO enrichment tab)). The 61 PPIs formed a network focused around the ergosterol synthetic pathway and specifically lanosterol 14-alpha-demethylase (Erg11), the direct target of fluconazole, one of four drugs we tested as environmental perturbation as phenotypes. We also tested 5-fluorocytosine (5-FC), a prodrug metabolized to 5-fluorouricil (5-FU) through the pyrimidine salvage pathway, which when incorporated into mRNA, disrupts protein synthesis and disrupts nucleotide synthesis through inhibition of thymidylate synthetase; metformin, a drug with no known target and a first-line treatment for type 2 diabetes; and trifluoperazine, an antipsychotic drug to treat schizophrenia that targets dopamine receptors in humans and calmodulins in yeast. In all four cases, the network of 61 PPIs act as “antennas” that, due to the small-world properties of PPI networks,^33^ probe changes in the biochemical network of the cell due to genetic variation.

We measured ∼430,000 PPIs (61 PPI × 354 strains × 5 conditions × 2 timepoints × 2 replicates) and thus the scalability of mrDHFR PCA was important to our devising a “pooled” screening strategy in which we introduced a chromosomal barcode to uniquely identify each of the 354 strains and integrated complementary mrDHFR PCA into the genome as fusions to the protein coding sequences of PPI binding partners (Figure 1C(i-ii) and Table S1). For chromosomal barcoding, donor DNA (Figure S2D and E) contained two NNNNN sites that uniquely label each of the 354 yeast strains (Table S2) in tandem with a *URA3* gene selection marker was integrated into the neutral locus YBR209W (Figure S2B and C).^34^ We then integrated the complementary mrDHFR PCA fragments into the genome, 3’ to one or the other of the open reading frames (ORF) of a protein pair, using a combination of CRISPR/Cas9 DNA cleavage with ORF-specific sgRNAs and homologous recombination (Figure S2A, D, and E). Thus, each pool consisted of all 354 barcoded strains with 1 of the 61 reporter mrDHFR PCAs integrated into their genomes. To serve as a baseline fitness corresponding to “zero PPI” in each pool (well), 2 strains (#43 and #599), were also barcoded but did not receive the mrDHFR fragments (Figure 1A(iii)). Throughout the barcoding of the strains and the tagging by mrDHFR fragments, all strains were maintained as haploid.

For the screen, we propagated the strains under MTX selection and in the presence or absence of one of the four drugs. Then, we performed genome extraction and PCR amplification of the barcode followed by NGS to estimate the fitness of each strain and, consequently, the magnitude of PPIs (Figure 1C(iii and iv)). To determine the genomic loci that significantly affect PPIs (henceforth called piQTL or protein-interaction quantitative trait locus), we performed linear regression between PPI strength and genetic diversity at each SNP locus (Figure 1C(v)).

Comparison of the mrDHFR-labeled strain’s growth in the presence of MTX (10 μg/ml) versus the untagged reference strain was used to determine PPI strength under “no drug” conditions (minimal media + supplements + MTX) and “with drugs” (drugs + MTX). The concentration of the drugs were determined in a prior GWAS study^26^ and PCA screen^11^ as 5-FC (0.2 μM), fluconazole (100 μM), metformin (50 mM), and trifluoperazine (17.5 μM). To estimate the growth rate of each strain, we extracted the genomic DNA of the pooled samples before and after selection, PCR amplified the locus containing the strain barcode, and then performed NGS (Figure 1c(iv)). The PPI strength of a protein pair (π) for an inbred strain (*i*) was calculated as the log-fold-change in barcode count (*n*) before and after selection, normalized by the response of the reference (untagged) strains: 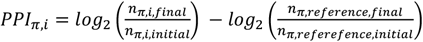. PPI strength estimates were strongly reproducible between two biological replicates (Pearson r = 0.88, *P*-value < 10^−16^; Figure 1G). Furthermore, the within-well controls also were highly correlated between the internal duplicates (Pearson r = 0.89, *P*-value < 10^−16^; Figure S3C), thus, we had sufficient accuracy for the PPI strength from the pooled measurements. We also recovered an average of ∼330 strains out of the 354 (∼93% of barcoded inbred strains) across all PPIs and drug conditions indicating sufficient power to perform QTL mapping (Figure 1F and Figure S1C).

Principal component analysis across all the conditions determined the largest source of PPI variation with each point representing the estimated change in PPI strength, averaged over all strains (Figure S3A,B). Expectedly, since mrDHFR tagging confers resistance to MTX, ∼65% of the variation (principal components 1 and 2) was due to MTX treatment (Figure S3A). In addition, we found that the drug conditions were largely overlapping, except for fluconazole (Figure S3B), which is expected, since the main functional target of the drug, Erg11 and other genes of the ergosterol pathway, were included in our set of 61 PPI reporters (Figure 1B).

### piQTLs occur in diverse genomic elements

We performed SNP to PPI association mapping analysis by linear regression at each SNP locus using the model 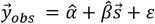, where 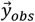 is the vector of measured PPI strength across all strains,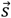 is the genotype, 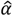 is the intercept, 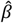 is the slope (the “effect size”) and *ε* is a random variable from an independent normal distribution.^35^ Since the inbred cohort have minimal linkage disequilibrium (Figure 1E), accounting for LD in our piQTL mapping analyses had negligible impact on our results (Figure S3D). We provide all association mapping results in a webserver (Figure S4) and list significant piQTLs in Tables S3-S6. Overall, we observed a total of ∼2,100 piQTLs (false-discovery rate (FDR) < 0.05) for all 61 PPIs under the 5 conditions (“no drug” and 4 drugs). These piQTLs were found in 473 unique SNPs. The largest number of piQTLs occur for fluconazole (Figure 2A; see also Figure S5A, where only the most significant SNP is considered within an LD block of r^2^ > 0.75, a typical cut-off ^36^). The piQTLs for all conditions were broadly distributed across the yeast genome (Figure 2A and Figure S5A). We expectedly found that the largest number of piQTLs are in protein-coding regions (Figure 2A and Figure S5A). Since the yeast genome is more compact than other model organisms and humans, and thus ∼6.3K out of the ∼12K SNPs in our strains are found in protein coding sequences (Figure S1B). Normalizing for the relative abundance of each genomic feature, ∼8% of these protein-coding SNPs are piQTLs under fluconazole, and ∼3% for the other drugs (Figure 2D). Surprisingly, within protein coding regions, piQTLs that are synonymous SNPs have comparable effect sizes to non-synonymous SNPs, except under fluconazole, where synonymous SNP effects were in fact slightly larger than non-synonymous effects (Figure S5D). It is possible that the elevated effect sizes for synonymous SNPs is due to linkage with nearby non-synonymous SNPs; however, this is unlikely because of the minimal LD in our inbred yeast cohort. Altogether, this result implies that the non-neutral effects of synonymous mutations on gene expression,^37^ manifests at the level of PPIs.

**Figure 2.**
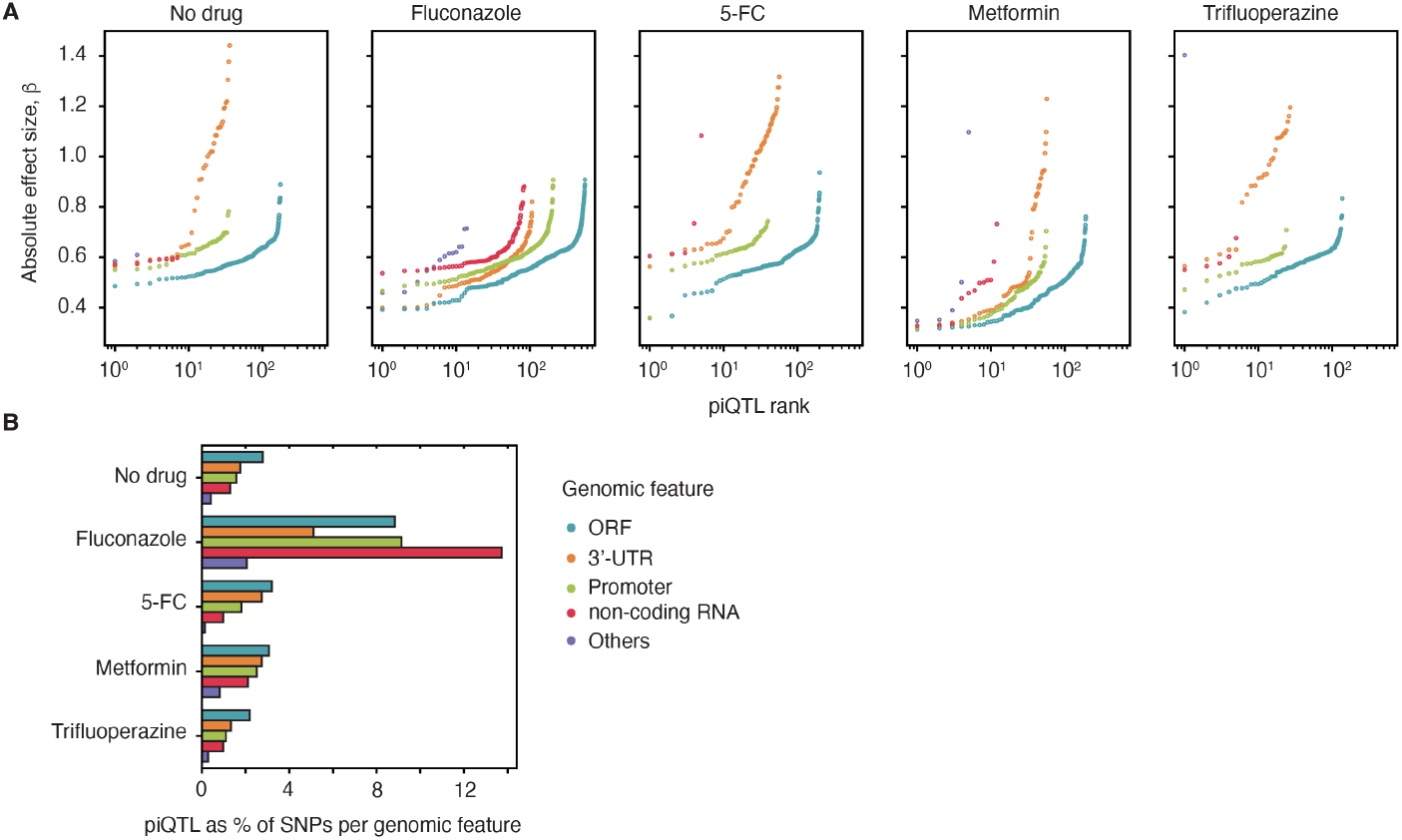
Genomic landscape of piQTLs. **(A)** Absolute effect sizes of significant piQTLs (FDR < 0.05) and their genomic locations in cells grown with no drug or treated with one of 4 drugs. The total number of piQTLs for each condition were 263, 974, 314, 332, and 201, respectively. **(B)** Proportion of SNPs that are piQTLs for each genomic feature (color legend is similar to panel A). See also Figures S4 and S5.

Most interestingly, SNP-to-PPI mapping also captured effects of genetic variations in non-coding RNAs, whose functions are not known or even annotated, including SUT (stable unannotated transcripts), CUT (cryptic unstable transcripts), and XUT (XRN1-sensitive unannotated transcripts)^38,39^ (Figure 2A). Although fewer piQTLs are found in non-coding RNAs than in 3’ UTRs or protein CDS, the magnitude of effect sizes are comparable under some drug conditions (Figure 2A, particularly panels for Fluconazole and 5-FC).

### Trans-acting are more abundant and have larger effects than cis-acting piQTLs

Analogous to cis- and trans-eQTLs, we can also classify the piQTLs based on their proximity to the genomic loci of the mrDHFR tagged interacting genes (Figure 3A). We defined cis-piQTLs as SNPs within 1 kb upstream or 200 bp downstream of either of the two interacting protein-coding ORFs, distance criteria utilized by a previous yeast genome-wide eQTL study.^40^ Conversely, trans-piQTLs are SNPs that are distant from the interacting proteins’ genomic loci. For instance, Miami plot comparing the piQTLs of the protein pair Erg11 (lanosterol 14-alpha-demethylase in chromosome 8) and Pis1 (phosphatidylinositol synthase in chromosome 16), with, and without fluconazole, shows a cis-piQTL in the intergenic region upstream of *ERG11*, only as a response to treatment with fluconazole (Figure 3B,C). This piQTL was also observed in a previous GWAS under fluconazole that used all 1,280 inbred strains from which we derived our subset of 354 strains, resulting in higher association power for the GWAS (Figure 3C).^26^ That piQTL can recover GWAS hits with a smaller number of strains suggests its power in detecting potential functional effects of a SNP locus to G×E.

**Figure 3.**
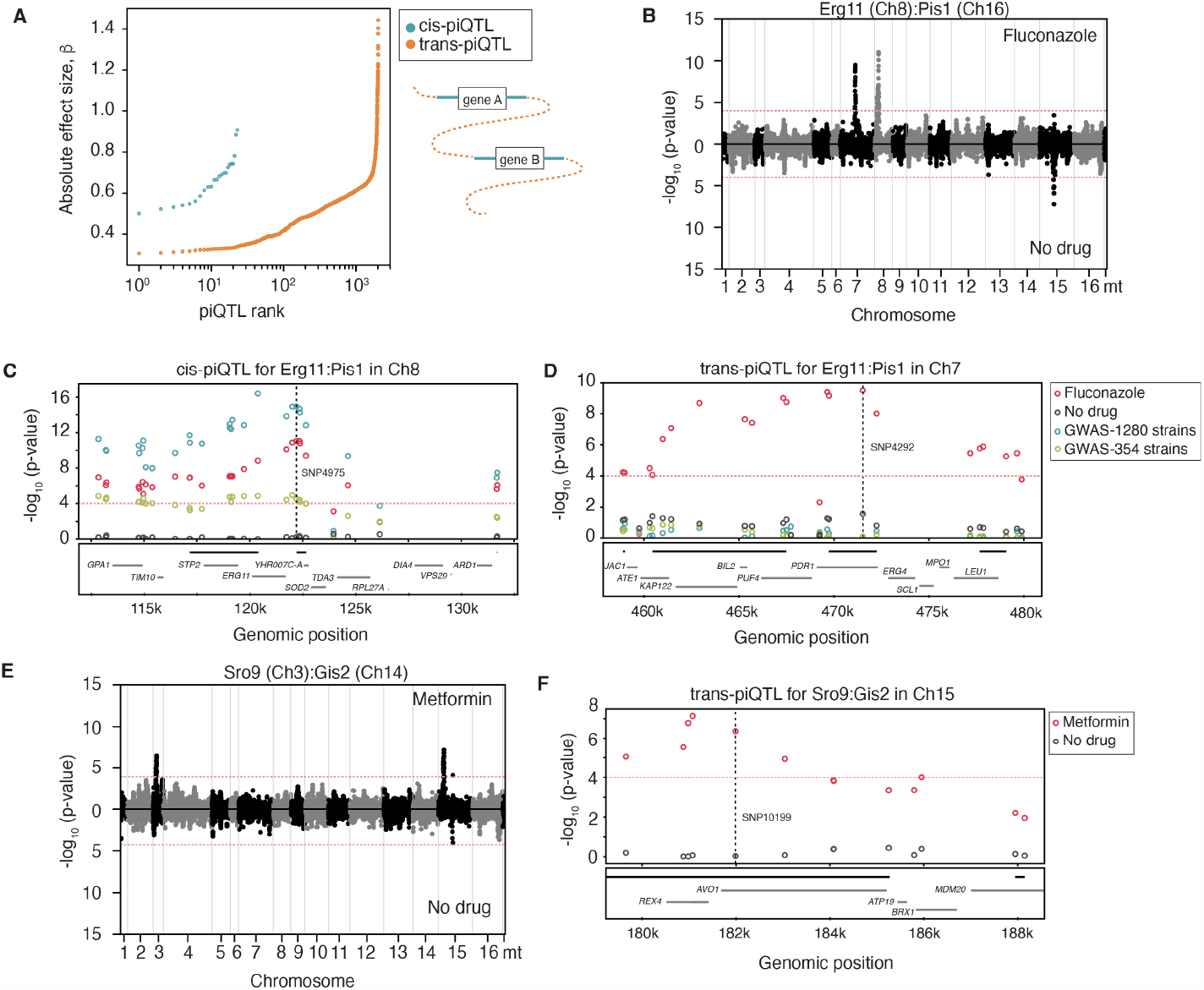
piQTLs local (in *cis*) and distant (in *trans*) to the gene loci of the interacting protein pair. **(A)** Effect sizes of cis- and trans-piQTLs in all conditions. Trans-piQTL outnumber and have greater effect sizes than cis-piQTLs (KS-test *P*-value < 10^−11^). See Figure S5A for cis- and trans-piQTLs per condition. **(B)** Comparison of the association for the PPI protein pair Erg11 in chromosome 8 and Pis1 in chromosome 16 across all ∼12K SNPs. **(C)** Zoom-in of the peak in chromosome 8 of panel B, which is a cis-piQTL near the *ERG11* gene locus. Erg11 is the target of Fluconazole. We also compared piQTL to a previous GWAS^26^ study using a complete 1,280 set of strains or the same 354 strains used in our study. See panel D legend. **(D)** Zoom-in of the cis-piQTL in chromosome 7 of panel D, which was undetected for fluconazole in the GWAS^26^ but a hit for other azoles. (**E** and **F**) Association plot for the PPI pair Sro9 in chromosome 3 and Gis2 in chromosome 14 in the presence or absence of metformin. Zoom-in of the cis-piQTL in chromosome 15 (panel F).*Note*: We follow the convention of writing gene names in italicized uppercase letters, while their protein products are non-italicized and only the first letter is uppercase.See also Figures S4 and S5.

A second peak in chromosome 7 centered around the gene *PDR1* (Pleiotropic Drug Resistance) is an example of a trans-effect (Figure 3B and D). *PDR1* encodes a transcription factor that regulates the overproduction of organic molecule transporters,^41^ conferring pleiotropic drug resistance, including to fluconazole in pathogenic fungal species.^42^ One of these is Hxt11, a transporter of a broad range of sugars and other molecules^43^ and a second, uncharacterized sugar transporter *HXT8;* both genes themselves have piQTLs (Table S3). *PDR1* is also adjacent to *ERG4* (coding for sterol C-24 reductase), the penultimate enzyme in the ergosterol pathway that is targeted by fluconazole (Figure 3D).

A unique and distinguishing feature of our results compared to eQTL and pQTL is that we observed more and stronger effect sizes for trans-than cis-acting piQTLs (Figure 2A),^8-10,20,44^ which holds true under all drug conditions (Figure S5B). This definition is robust to our cut-off distance of cis- and trans-acting SNPs since the majority of trans-piQTLs are inter-chromosomal (Figure S5C, Tables S3 to S6). Even when cis-acting SNPs are defined to be within 50 kb of the protein pair loci, ∼88% of the piQTLs are still classified as trans-acting (Figure S5C). This observation supports the view that PPIs captures the entire fate of a gene, including variations in passive processes that effect the abundance of proteins including transcription, translation and protein degradation, and active processes that affect strengths of PPIs, such as post-translational modifications (Figure 1A). Detecting genetic interaction via mRNA or protein abundance is performed indirectly by correlating these molecular readouts between two genes; in contrast, PPI is a direct measure of the physical interaction of two genes’ protein products. An important consequence of these results is that using PPIs as reporters of genetic variation effects could help define the molecular mechanisms underlying the epistasis and pleiotropy of complex traits.

### piQTLs reflect the topology of biochemical networks

At the outset, we hypothesized that because of the small world nature of the PPI network, assessing effects of genetic variation on the response of the cell to environmental perturbation may only require a few reporters. Consistent with this hypothesis, ordering the 61 PPIs according to their proximity on the PPI network (from Figure 1B and Figure S7E), the PPIs that are closely connected in the PPI network, particularly those sharing the same protein partner, are more likely to share piQTLs (Figure 4A). We also observed that the number of overlapping piQTLs between pairs of PPIs increase as their distance on the PPI network decreases (Figure 4B). Despite the broad genomic distribution of piQTLs in the genome, they were found in genes coding for proteins that are close to each other on the PPI network. For example, we observed high connectivity and shorter distance, compared to random, between proteins containing piQTLs for Erg11:Orm2, as shown on the sub-network of the *in vivo* protein interactome (Figure 4C and D),^11^ which is true for other reporter PPIs (Figure 4G). However, we observed the opposite results when the Erg11:Orm2 piQTLs are projected onto the genetic interaction network (GIN) defined by comprehensive double-gene knockouts,^45^ where the piQTLs are more spread over the network (Figure 4E and F). This trend holds true for piQTLs of other PPIs under fluconazole (Figure 4G), as well as other drug conditions (Figure S6). We also observed that piQTLs are not enriched *within* functional modules defined by gene-knockouts, rather piQTLs are enriched in the intervening nodes *between* the modules (Figure 4F and Figure S6). Based on these results, we hypothesize that the gene knockouts reflect drastic effects on PPIs, in contrast to effects of polymorphisms that are part of the standing genetic variation, such as the ∼12K SNPs between the two parental strains (Figure 1D). Thus, PPI-SNP mapping provides a unique measurement of the sensitivity of the PPI network to standing genetic variation, which is complementary to gene knockouts or random mutagenesis by CRISPR/Cas9.

**Figure 4.**
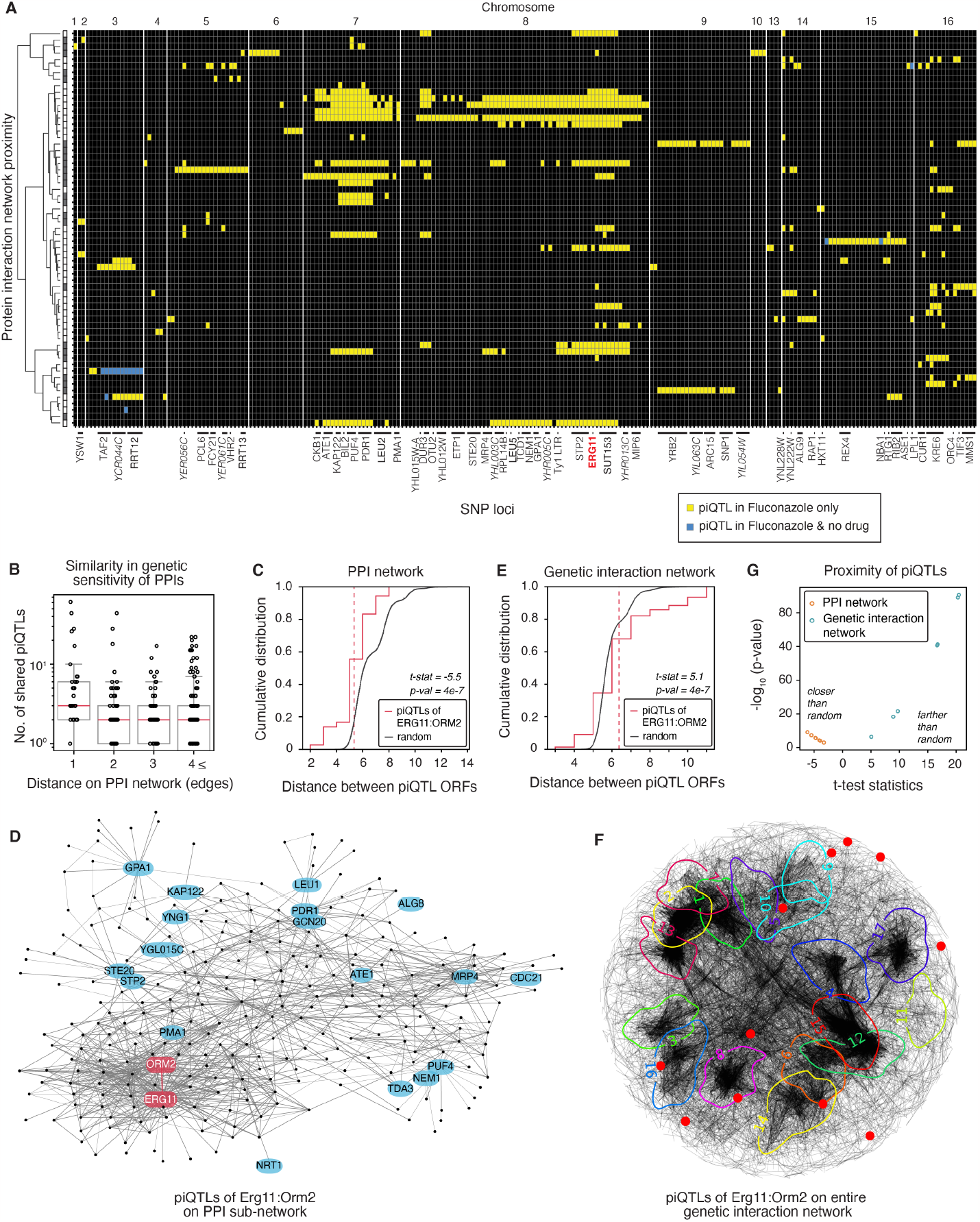
piQTLs reflect the topology and strength of biological networks. **(A)** SNPs that were piQTLs in at least one of the 61 PPI reporters. Yellow boxes are significant piQTLs identified only upon fluconazole treatment. Blue are piQTLs in both with and without fluconazole treatment, but whose FDR changed by an order of magnitude with the drug. piQTLs that have comparable P-values in both conditions are not shown but listed in Table S3. The rows are ordered according to the hierarchical clustering of the 61 PPI probes based on their distance on the PPI network (see Figure S7E for PPI identities). **(B)** Similarity in genome-wide association between two PPIs based on the number of overlapping piQTLs versus their distance on the PPI network. **(C** and **D)** piQTLs tend to cluster in PPI networks compared to random as quantified by their pairwise distances (panel C) and demonstrated by an example (panel D). **(E** and **F)** piQTLs under fluconazole were projected onto the genetic interaction network defined from double knockouts in yeast.^45^ piQTLs tend to be between distant pairs compared to random and do not cluster within functional modules defined by the knockouts, as shown by an example (panel F). See Figure S6 for the definition of the modules and projection of piQTLs on the GIN for other drug conditions. **(G)** Generalization of panels C & E to other PPIs with more than 5 piQTLs (panel A). See also Figures S6 and S7.

### non-coding RNAs affect protein-protein interactions

The surprising abundance of piQTLs in non-coding RNA (ncRNA) regions (Figure 2A) prompted us to determine how these effects partition into the different classes of non-coding RNAs (Figure 5A). In *S. cerevisiae*, there are ∼2, 000 ncRNAs that make up ∼20% of its genome.^46^ Two classes of ncRNAs were initially identified according to their half-life in the cell, the stable unannotated transcripts (SUTs) with relatively long half-lives and cryptic unstable transcripts (CUTs) RNAs with short half-lives.^38,46^ Deletion of the cytoplasmic exonuclease Xrn1, followed by RNA sequencing, revealed another class of ncRNAs termed Xrn1-sensitive unstable transcripts (XUTs),^39^ some of which overlap with either SUT or CUT. Since, Xrn1 occurs only in the cytoplasm, SUTs or CUTs that are also XUTs are exported to the cytoplasm and processed like regular mRNAs.^39^ The full mechanism and role of these transcripts are generally unknown and a subject of debate.^47^

**Figure 5.**
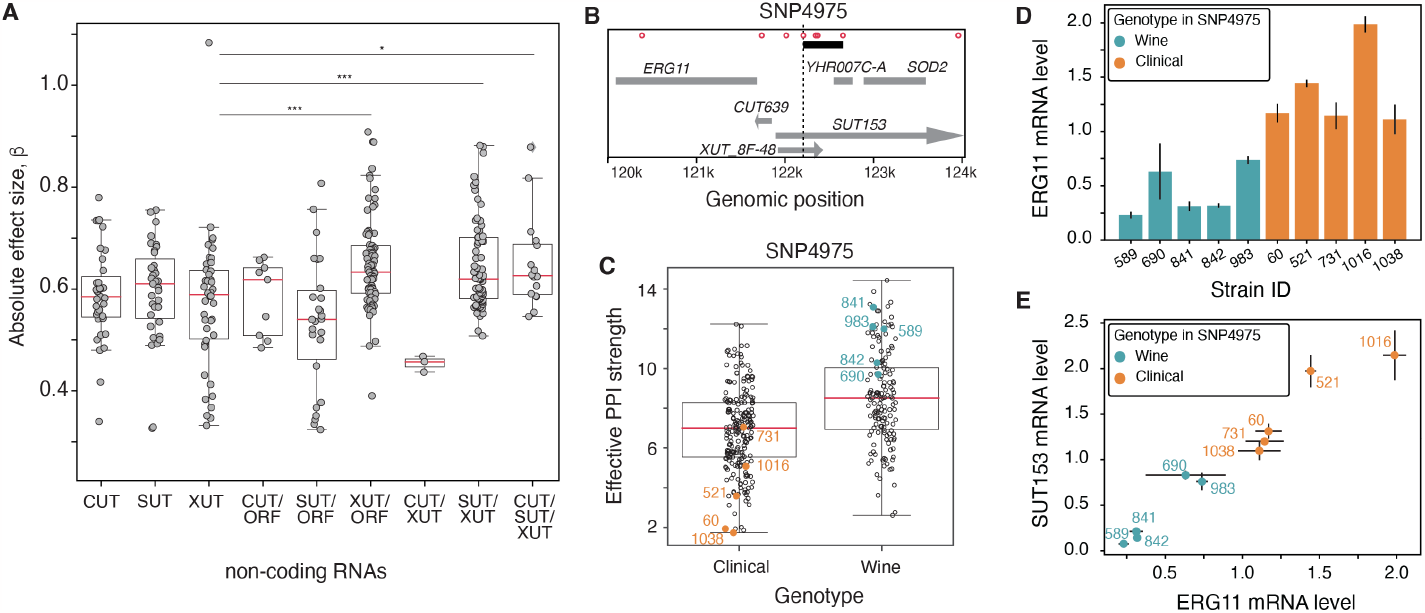
Non-coding RNAs significantly affect protein-protein interactions. **(A)** Effect sizes of the piQTLs in non-coding RNAs CUT, SUT, and XUT. Grouped separately are SNPs that have multiple annotations, either overlapping with ORFs or other non-coding RNAs. **(B)** Zoom-in of the genomic locus near SNP4975 showing the most significant piQTL in fluconazole-treated cells near *ERG11*. This region contains multiple overlapping non-coding RNAs. Red dots indicate all SNPs in the region (see also Figure 3C). Black line indicates an LD block with *r*^*2*^ > 0.75. **(C)** PPIs of strains grouped according to their genotype at the SNP4975 locus. **(D** and **E)** qPCR quantification the transcripts for *ERG11* and *SUT153* using five strains from each of the parental wine and clinical isolate genotypic backgrounds (colored dots in panel C). mRNA level is expressed relative to the *ACT1* gene (Actin).

We found that the effect sizes vary across all types of non-coding RNAs (Figure 5A). Interestingly, the piQTLs in non-coding RNAs that are multiply annotated have higher effect sizes, particularly the XUTs that are exported to the cytoplasm and processed as regular mRNA transcripts and are thus, more likely to be functional (Figure 5A). This could indicate that these loci either have dual effects, via their product proteins or through the non-coding RNA itself. One of the strongest effect sizes for fluconazole was SNP4975 in the ∼2kb-long *SUT153*, which is also annotated to overlap with the shorter ∼0.5 kb *XUT_8F-48* (shown for the PPI Erg11:Pis1 in Figure 5B and Figure 3C). This locus also showed a significant peak for 16 other PPIs (Figure 4A) and is the only significant locus around *ERG11* for 4 of these PPIs, suggesting that it is the likely causal SNP in the region. Thus, we could hypothesize that *SUT153* regulates the transcription of *ERG11* and its intracellular abundance and interaction with other proteins. In particular, a smaller fragment of *SUT153*, which is *XUT_8F-48*, is found in the cytoplasm (Figure 5B)^39^ and does not overlap with the adjacent gene (YHR007C-A, a putative protein of unknown function). SNP4975 is a candidate cis-acting piQTL, affecting a gene coding for one of the interacting pair of proteins that we probed. To test our hypothesis, we quantified the mRNA level of *ERG11* and *SUT153* by qPCR for 10 strains, half containing the SNP from the clinical parent and the other from the wine parent (Figure 5C).

We found that the two groups of strains differed in their *ERG11* expression, reflecting the strong sensitivity of this locus to fluconazole (Figure 5D). Moreover, the *ERG11* and *SUT153* mRNA levels were strongly correlated (Figure 5E), suggesting a possible active regulatory role for this ncRNA. The next most significant ncRNA in our screen was *SUT093* (also *XUT5R_135*) in Chromosome 5, which is directly upstream the gene *VHR2* (regulator of the Vitamin H transporter Vht1) that is required for the expression of ergosterol pathway enzymes Erg6 and Erg9 ^48^ (Table S3). Altogether, this finding highlights the PPI network’s ability to probe genomic elements whose function are unannotated or unknown.

### piQTLs recover known and novel drug effects

Having established that the small world nature of PPI networks and the sensitivity of PPI to genetic variation can report on broad cellular processes (Figure 1A), we next determined if the piQTLs also recover known drug mechanisms and could provide hypotheses for new ones. To prioritize the piQTLs for inferring drug mechanisms, we performed a meta-analysis to combine the significance of a SNP as a piQTL across *all* 61 reporter PPIs into one statistical score. Meta-analysis calculates an average effect size (and P-value), weighted by the precision of each 61 SNP-to-PPI linear regressions (Methods) (P-values are reported in Tables S3 to S6). We found that SNPs that survived the FDR cut-off for piQTL in multiple PPIs have higher significance in the meta-analysis (Pearson correlation *P*-value < 10^−16^; see Figure S7F and full results in Tables S3 to S6). This analysis also simplified the task of prioritizing piQTLs and their corresponding genes for inferring mechanisms of action of the four drugs we tested.

To determine if our piQTLs are consistent with known mechanisms of action of drugs in the literature, we queried PubMed for studies that indicated the role of a piQTL-containing gene to the drug. For instance, for fluconazole we searched it and other azoles, as well as the drug’s ability to inhibit the fungal cytochrome P450.^49^ We found that the number of publications relating a gene to fluconazole is higher for genes containing the most significant piQTLs (Figure 6A, Pearson correlation *P*-value <10^−2^), which is not the case for the “no drug” condition (Pearson correlation *P*-value is not significant).

**Figure 6.**
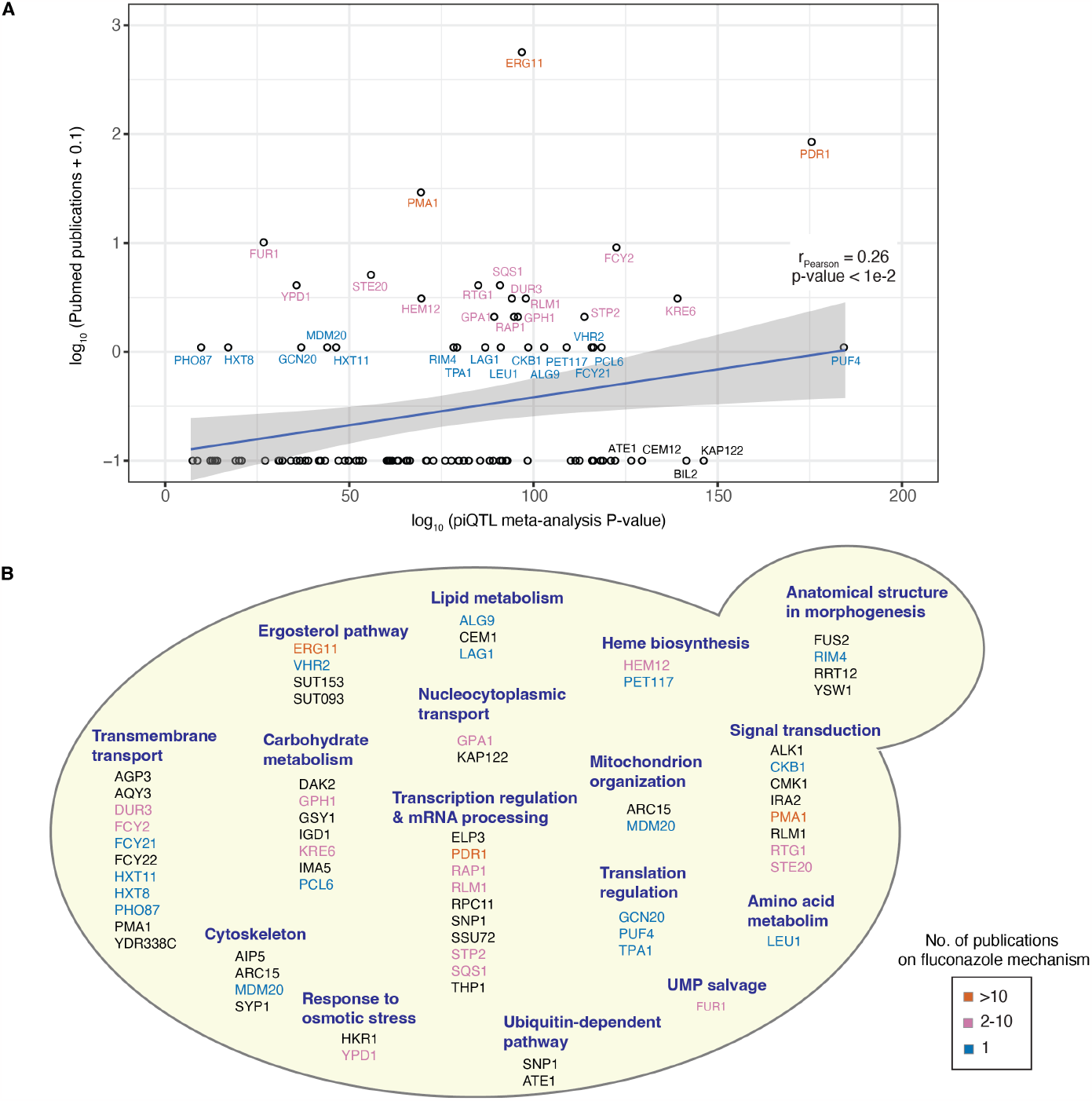
piQTLs in fluconazole and reported mechanisms of fungal resistance. **(A)** Significance of piQTLs within a gene and the number of studies in PubMed relating that gene to mechanisms of fluconazole, other azoles, and fungal resistance. Colors for number of publications correspond to panel a legend. Gene descriptions are listed in Table S3. **(B)** Biological processes of significant piQTLs under fluconazole.

Despite having only 61 reporters centered around the ergosterol pathway, piQTL recovers broad biological processes and identifies new crucial players (Figure 6B). As highlighted above, a primary major mode of resistance for fluconazole is the overexpression of transporters, particularly those whose genes are regulated by the transcription factor Pdr1 (Figure 3D). However, we also found other putative transporters whose function is entirely unknown (YDR388C and *AQP3*) or known transporters but whose role in azole resistance is not yet established (*AGP3*) (Figure 6B and Table S3). A well represented set of genes are kinases involved in signal transduction as a response to cellular stress (Figure 6B), a logical result since antifungals induce stress signal transduction pathways.^50^

Fungal membrane homeostasis requires the concerted production and maintenance of sterols, as such, we found piQTLs in *ERG11* (Figure 4A; Figure 6B) and upstream non-coding RNAs that could potentially regulate *ERG11* expression and protein abundance (Figure 5B,E). Sterol biosynthesis is strongly coupled to the production of other components of the plasma membrane, such as sphingolipids, which are important components of lipids rafts.^51^ Cellular stress due to azoles induce the expression of the gene *LAG1* (longevity assurance gene 1), which codes protein involved in the synthesis of sphingolipid ceramide, a regulator of azole resistance in *C. albicans*.^51,52^ Other components of the lipid rafts that are also piQTL is the plasma membrane proton pump Pma1 (Figure 6B).

Compared to upregulation of ERG genes to rescue lipid membrane integrity and the expression of transporters to reduce cellular drug concentration, a less explored mechanism of antifungal tolerance is the role of respiration and mitochondrial organization. Fungal defects in mitochondrial respiration leads to “petite”, slow-growing, mutants that are resistant to antifungals.^53^ A piQTL in *PET117* (gene name “petite” colonies 117), whose deletion leads to the “petite” phenotype,^54^ has only recently been shown to couple heme synthesis to cytochrome oxidase (Figure 6B).^55^ Another piQTL is in *LEU5*, although not yet directly implicated in fluconazole, is a yeast mitochondrial carrier required for the accumulation of Coenzyme A in the mitochondrial matrix.^56^ Lastly, we detected a piQTL in *RTG1*, a transcription factor for genes that code for proteins involved in mitochondrion-to-nucleus signaling that has been shown to upregulate expression of multi-drug resistance gene *PDR1*, which also contains a piQTL (Figure 3D).^57^ These piQTLs provide hypotheses and candidate genes that mediate the function of mitochondria in fungal drug resistance.

Interestingly, the purine-cytosine permease *FCY2* that is the target of 5-FC, including two other putative homologs *FCY21* and *FCY22*, are also piQTLs under fluconazole, suggesting a mechanism of cross-resistance between the two antifungals.^58^ Contrary to our expectations, these resistance conferring genes for 5-FC do not appear as piQTLs for 5-FC, which could indicate that the 5-FC concentration used in the study may be low to elicit PPI dependence on these loci (Figure S7A; Table S4). Broadly, these results highlight the complexity and pleiotropy of G×E, even for drugs with well-defined targets.

Despite the small number of 61 PPI reporters and the paucity of connection between metformin and ergosterol pathway, which is enriched among the PPI reporters, we were able to capture known mechanisms surrounding metformin in humans (Figure 7). For example, metformin exhibits direct inhibitory effects on mTOR (mammalian target of rapamycin), a pivotal nutrient-sensitive kinase that controls growth and cell proliferation in response to glucose, energy, growth factors and amino acids. The piQTL in *AVO1* (*a*dheres *vo*raciously to Tor2 protein 1) (Figure 3F) is directly part of the mTOR complex ^12,60,61^. While another piQTL in the G-protein *ASC1*, whose human homolog *RACK1* (receptor of activated protein C kinase 1), inhibits mTORC1 and is regulated by rapamycin through changes in both mRNA and protein levels.^62^ Furthermore, our previous screen showed that *ASC1*’s ability to form a homo-dimer complex is sensitive to rapamycin treatment.^12^

**Figure 7.**
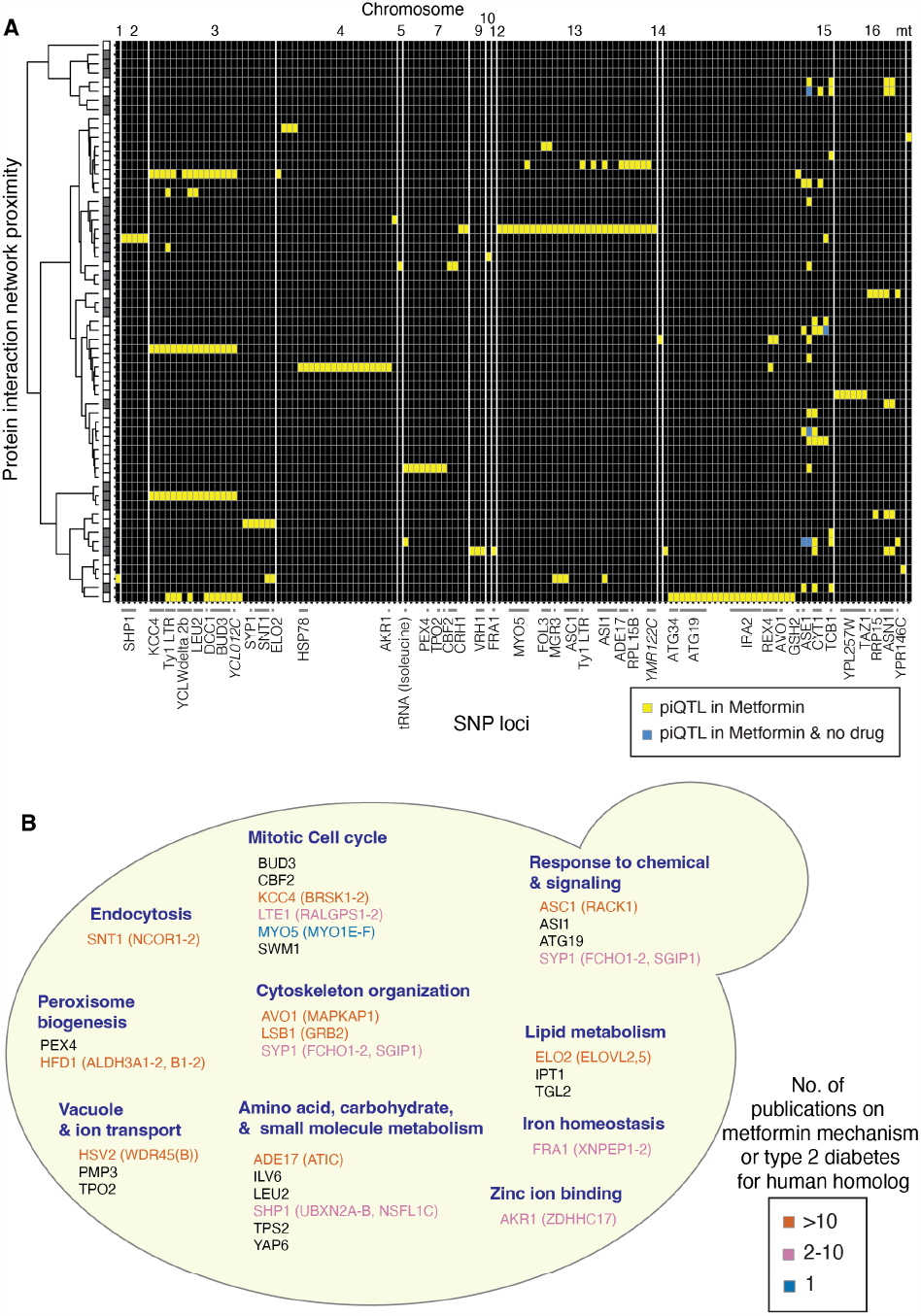
Metformin piQTLs predict roles of zinc and peroxisomes in cytoprotection and ferroptosis. **(A)** Genome-wide piQTL across all 61 PPI under metformin. The rows are ordered according to the hierarchical clustering of the 61 PPI probes based on their distance on the PPI network (see Figure S7E for PPI identities). **(B)** Yeast piQTLs and their human homologs in parenthesis are colored according to number of publications implicating the gene or its human homologs to metformin or diabetes. piQTLs associated endocytosis, vacuole function, including and peroxisome biogenesis are implicated in ferroptosis, which can be induced by metformin in cancer persisters cells by increasing free iron.^59^ Gene descriptions are in Table S5.See also Figure S7.

piQTLs also occurred in yeast genes whose human homologs are strongly implicated in type 2 diabetes (Figure 7B and Table S5), although the direct connection to metformin is not yet understood. These include a putative subtilase-type proteinase *RRT12*’s whose human homolog, *PCSK9* (proprotein convertase subtilisin/kexin type 9), is a crucial player in cholesterol homeostasis and low-density lipoprotein (LDL) catabolism.^63^ Indeed, *PCSK9* is a target for drugs to lower LDL^64^ and its expression is reduced by metformin in humans and mouse models of diabetes.^65,66^ Another piQTL occurs in fatty aldehyde dehydrogenase *HFD1* dehydrogenase involved in ubiquinone and sphingolipid metabolism, and its human homologs aldehyde dehydrogenases *ALDH3B1* and *ALDH1A1* have been associated with type 2 diabetes and other metabolic disorders in GWAS studies,^67^ particularly *ALDH1A1* which is differentially expressed in disease models of diabetes and obesity.^68^

Our results also provide novel hypothesis on some perplexing observations regarding metformin. We recently discovered that metformin causes dissociation of iron from proteins in yeast,^12^ likely by competing for iron binding to heme, to iron-sulfur clusters, and to wide-range of metabolic activities requiring iron.^69^ Indeed here, we also found significant piQTLs among iron homeostasis genes such as *FRA1* that controls iron regulon transcription (Figure 7B).^70,71^ A consequence of this shift in iron binding is that metformin can induce cell death by ferroptosis in cancer persisters cells, which can escape drug treatments.^59^ This is an important observation because other ferroptosis-inducing compounds that target cancer persisters also kill normal cells that are susceptible to ferroptosis, including neurons and cardiomyocytes.^72,73^ Metformin, however, has no such affects and in fact, could be neuro-protective.^74,75^ Thus, any insights into the differences between modes of ferroptosis induction by metformin versus other drugs could be valuable in developing new strategies to specifically kill only cancer persisters. We found several significant piQTLs among genes associated with cellular sub-compartments implicated in ferroptosis (Figure 7B); specifically, the axes of iron uptake and extraction begin with endocytosis of iron binding proteins, uptake into lysozomes (vacuole), degradation of the proteins and transport of iron out of the lysozomes. A surprising result is that, while piQTLs were observed in some genes encoding proteins associated with iron, all the vacuole metal transporters are for other metals, notably zinc. Zinc has been implicated in improving lipid metabolism and renal function in diabetes patients who take metformin and neuroprotection in animal models.^76^ This result raises the interesting hypothesis that metformin may fine-tune a balance between protective levels of zinc versus toxicity of iron and other metals such that it can induce or protect against ferroptosis in different contexts.

We also observed piQTLs in a key set of genes implicated in peroxisome biogenesis (Figure 7B). Peroxisomes have only recently been implicated in ferroptosis, in a genome-wide CRISPR-Cas9 suppressor screens of cancer persister cells.^77^ Furthermore Zou et al.^77^ showed that peroxisomes contribute to ferroptosis by synthesizing polyunsaturated ether phospholipids (ePL), which as substrates for lipid peroxidation, results in ferroptosis. Neurons and cardiomyocytes have high levels of ePLs, and, as noted above, which could explain why they are susceptible to ferroptosis.^78^

Finally, the antipsychotic trifluoperazine, primarily used to treat schizophrenia, binds to and acts as an antagonist to adrenergic, dopaminergic, and acetylcholine receptors. Notwithstanding the complexity of the mechanism of trifluoperazine in human patients, some of the yeast piQTLs are in genes whose human homologs are associated with schizophrenia (Figure S7C,D), including, *RACK1*, which exhibits abnormal localization in schizophrenic patients^79,80^ and interacts with dopamine receptors.^81,82^ Trifluoperazine also has strong anticancer activity, hypothesized to be due to mitochondrial stress associated with proteosome-mediated degradation,^83-85^ biological processes that are enriched in our piQTLs and their human homologs (Figure S7D).

## DISCUSSION

Here, we showed how PPI networks can be accurate reporters of the molecular and functional consequences of genomic variation. piQTL has several potential advantages over eQTL and pQTL. First, the broad applicability of pQTL faces practical limitations in that proteomics, typically done by mass spectrometry, is technically more challenging and costlier than transcriptomics by RNA-seq, which takes advantage of next-generation sequencing (NGS). Our implementation of pooled PPI screen shows that piQTL has the practicality and scalability of RNA-seq and eQTL because the PPI readout is through NGS. Second, piQTL captures the integrated effects of genetic variation on the proteome, thus a better reporter of G×E than eQTL or pQTL. Finally, piQTL could accurately report on multiple cellular processes that are close to phenotype, such as protein localization.

The preponderance of GWAS QTLs and eQTLs found across the genome and over many genes do not have an interpretable connection to the phenotype or disease of interest.^1,2^ This quandary led Pritchard *et al*. to propose the “omnigenic” model of complex traits.^10,86^ This model postulates that gene regulatory networks are sufficiently interconnected such that all genes expressed in trait-relevant cells are likely to affect the functions of core trait-related genes. We argue that such inter-connectivity is well-established in the small-word characteristics of PPI networks,^33^ which manifests in the abundance of trans-piQTLs over cis-piQTLs. Consequently, we postulate that the omnigenic nature of complex traits could best be explained by PPI networks.

Additionally, the complexity of genotype-phenotype-environment relationship (G×E) is due to non-additive interactions between mutations in different genomic loci (epistasis) and on the ability of a single gene locus to affect multiple phenotypes (pleiotropy).^4^ Since *in vivo* PPI directly measures the dynamic physical interactions of the gene’s protein products, it could also provide direct quantification of epistasis and pleiotropy. Taken together, our works provides a general roadmap to the discovery of molecular mechanisms of complex phenotypic traits via PPI networks.

### Limitations of the study

Our results support the hypothesis that PPIs are downstream reporters of the consequences of genetic variation on expression and protein abundance. However, to determine what is uniquely captured by PPI versus protein abundance or gene expression will require follow-up quantitative comparison and statistical colocalization analysis with pQTL and eQTL, ideally using the same strains and conditions used in this study. Additionally, how genetic perturbation percolates across these molecular phenotypes is also a function of drug concentration, which must also be explored in future studies. We anticipate that the effect sizes will not necessarily correlate linearly with the drug concentration, since from biochemical arguments, the number of PPI complexes, and binding curves in general, are sigmoidal with respect to drug concentration. Lastly, our findings, currently based on 61 *in vivo* reporters, will need to be generalized to a larger number of PPI interactions to capture biological processes insufficiently covered by our sub-network, but could be relevant to other drug conditions.

## ACKNOWLEDGEMENTS

The authors thank Dan F. Jarosz (Stanford University) for sharing the inbred strain collection. We also thank Jean-Christophe Grenier and Raphaël Poujol for technical input on the bioinformatics analyses. We acknowledge funding from the following agencies and foundations: Canadian Institute of Health Research grants MOP-G-408523 (AWRS) and MOP-GMX-152556 (SWM); Natural Sciences and Engineering Research Council of Canada grant RGPIN-2016-06566 (AWRS); Canada Research Chairs (SWM, AWRS); Human Frontiers Science Program grant RGP0034/2017 (SWM); Fond de Recherche du Quebec – Santé (FRQS) and IVADO (JGH); and Uehara Foundation of Japan (TS).

## AUTHOR CONTRIBUTIONS

Conceptualization: SB, TS, L Gauthier, SWM, AWRS; Methodology: SB, TS, L Gauthier, ZS, OP, L Gonzalez, XC-G, CM, NC, JGH; Investigation: SB, TS, L Gauthier, SWM, AWRS; Visualization: SB, TS, OP; Funding acquisition: SWM, AWRS; Project administration: SWM, AWRS; Supervision: SWM, AWRS;

Writing – original draft: SB, TS, SWM, AWRS; Writing – review & editing: SB, TS, L Gauthier, SWM, AWRS.

## DECLARATION OF INTERESTS

The authors declare no competing interests.

## STAR METHODS

## CONTACT FOR REAGENT AND RESOURCE SHARING

Further information and requests for resources and reagents should be directed to and will be fulfilled by the lead contact Adrian Serohijos.

## EXPERIMENTAL MODEL AND SUBJECT DETAILS

### Inbred strain collection

The population cohort in this study consisted of 354 haploid inbred strains of *S. cerevisiae* previously constructed and genotyped by She and Jarosz for a GWAS study.^26^ The strains were F6 progenies from inbreeding of two parental strains RM11 (wine isolate) and YJM975 (clinical isolate). There were 12,054 high-frequency SNPs in the parental and inbred strains. During our genomic editing to add the strain barcode and mrDHFR fragments, the cells were kept haploid using a mating inhibitory peptide.^87^

## METHOD DETAILS

### Annotation of the SNPs genomic features

To determine the genomic elements on which the 12,054 SNPs are located, we obtained the annotated reference genome of *S. cerevisiae* from the Saccharomyces Genome Database (SGD) (version R 64-3-1 released on 2021-04-27). The annotated features include all open-reading frames (ORFs), the 5’ and 3’ UTR of these ORFs, non-coding RNAs, intergenic regions, and other genomic features (long tandem repeats (LTRs), transposons, and telomeric regions). We also mapped the annotations of promoter regions and unstable transcripts performed by Rossi and co-workers.^88^ Reference datasets and results of the annotation mapping pipeline are available in *GitHub* (https://github.com/ladyson1806/Serohijoslab-piQTL).

### Subnetwork of 61 PPIs

The 61 PPIs (Figure 1B; Table S1) targeted in this study were selected from a previous proteome-wide *in vivo* interactome screen by mrDHFR-PCA in the *S. cerevisiae* S288C background.^11^ The PPIs were also curated to cover a wide range of molecular functions and biological processes (Table S1).

### Genome editing for chromosomal barcoding and DHFR fragment tagging

Our yeast editing approach for barcoding and DHFR tagging used a combination of homologous recombination for accuracy and CRISPR/Cas9 for efficiency (Figure S2A).^89^ Briefly, a “donor DNA” with homology arms to the target genomic locus, an sgRNA targeting the locus of integration, and a linearized Cas9 expression vector were co-transformed into yeast. Gap repair led to the reconstitution of the Cas9 expression plasmid, followed by the cutting of the genomic target locus by Cas9/sgRNA, and its subsequent repair by homologous recombination. The primers used in this section are summarized in Table S7.

#### (a) Competent cell preparation and transformation

Yeast-competent cells were prepared by following the lithium acetate protocol.^90^ The frozen stock of the yeast cells was incubated overnight at 30°C in YPD medium supplemented with 50 nM mating inhibitory peptide (SKGSPINTQYVF, BioBasic).^87^ The following day, the pre-cultured yeast cells were diluted in pre-warmed 2x YPD media to a concentration of 2×10^6^ cells/mL. Cells were incubated until OD_600_ of 0.8-1.0 (∼8×10^6^ cells/mL), at which point the cells were harvested by centrifugation. Cells were rinsed once using sterile water before being reconstituted in Frozen Competent Cell (FCC) solution (5% v/v glycerol, 10% v/v DMSO). The quantity of FCC solution utilized equated to 0.01 times the volume of the culture medium used. The cell suspension was dispensed into microcentrifuge tubes in 50 μL aliquots and gradually frozen at −80°C. For transformation, the frozen competent cells were thawed in a 37°C water bath for 15-30 seconds and briefly centrifuged to remove the supernatant. The cells were resuspended in 360 μL of transformation mix (10 μL salmon sperm DNA, 260 μL 50% w/v PEG 3350, 36 μL 1.0M LiOAc, transforming DNA, and fill-up with sterile water). The cell suspension was then heat-shocked at 42°C for 30 min and briefly centrifuged to remove the supernatant. The cells were resuspended in 1 mL YPD medium, incubated at 30°C for 4 h, and plated on appropriate selective YPD plates.

#### (b) Chromosomal barcoding of inbred strains

To perform the chromosomal barcoding, the donor DNA (Figure. S2B) contained two NNNNN sites that uniquely label each of the 354 yeast strains (Table S2) in tandem with a *URA3* gene selection marker that allows selection in minimal media without uracil. To determine the intrinsic reproducibility of fitness estimates within each pool, 3 strains (#17, #40, and #180) were labeled with two unique barcodes. Thus, the total number of barcodes is 357 corresponding to 354 unique inbred strains.

The complete barcode sequence consisted of two short barcode sequences placed on either side of a 20 bp consensus sequence. There were 24 unique sequences in each of the left and right short barcodes, resulting in 576 possible unique barcode combinations, of which we used 357 (Table S2). These barcodes were also designed to have comparable GC content and at least three Hamming distances from other barcodes. The strain barcodes were inserted into the YBR209W dubious and neutral open reading frame, whereby its disruption does not have effects on fitness.^34^

The *URA3*-barcode DNA cassettes were prepared by assembling the YBR209W left homology arm (175 bp, gcctgatattcagaagtgagcgagtatacaaatctcacacctcatctgctctagtgtttaaatacaaacattcaatttccaa caacgtctaacaaagctcgattcaaattaagaagcaggaaaaagcaaaacatctgcgtgtcttatctcaatggcttgcgctaactgcgaa cag), the *URA3* gene, left barcode oligo (gtgcgacagcttcagacctgNNNNNgcccaatacgcaaaccgcct), right barcode oligo (cgcatctttggctgtatttcNNNNNgtgcgacagcttcagacctg), and the *YBR209W* right homology arm (175 bp, gaaatacagccaaagatgcgtgccgttgcagttagctaacaacctggcgttcggcgatcgccataaga gatctgccaattttaaaaagggtattgccaatacacatagtagccttatctgcagcaagccataggactaatgtgttcgacgtcgttggggaa). The YBR209W homology arms were amplified by PCR directly from the inbred strain genome, and the *URA3* cassette was amplified by PCR from plasmid pWS158 (Addgene plasmid #90517). The amplified DNA fragments, barcode oligos, and YBR209W left-/right-homology arms were assembled together by fusion PCR.

DNA barcode labeling of the yeast library strains was performed in accordance with the CRISPR yeast genome editing protocol (Figure S2A).^89^ To prepare the linearized sgRNA expression cassette for targeting the YBR209W locus, oligos containing the CRISPR sequence of YBR209W were inserted into pWS082 (Addgene plasmid #90516), the template plasmid of sgRNA cassette, using BsmbI Golden Gate Assembly. The linearized sgRNA expression cassette targeting YBR209W was then amplified from pWS082-YBR209W using PCR. The linearized Cas9 cassette was amplified from pWS176 (Addgene plasmid #90963) by PCR, and the donor DNA was the *URA3*-barcode cassette. The three cassettes were gel purified by extracting fragments of the following sizes: sgRNA cassette, 1045 bp; Cas9 cassette, 9993 bp; URA3-barcode cassette, 1488 bp. The purified DNA cassettes were mixed in the following ratio and transformed into competent cells: sgRNA cassette, 200 ng; Cas9 cassette, 100 ng; and URA3-barcode cassette, 5 μg. The transformed cells were stored at −80°C for downstream processing. Barcode insertion was validated by plating transformed cells, and then colony PCR and Sanger sequencing of 16 randomly picked colonies (Figure S2C).

#### (c) mrDHFR-fragment genomic tagging in yeast barcoded library strains

Prior to mrDHFR tagging, we pooled the barcoded inbred strains at equal densities and then performed mrDHFR fragments tagging via CRISPR yeast genome editing protocol (Figure S2A). To prepare the linearized sgRNA expression cassettes for targeting loci, oligos containing the CRISPR sequence (table S1) were inserted into pWS082 (Addgene plasmid #90516), the template plasmid of sgRNA cassette, using BsmbI Golden Gate Assembly. sgRNAs were designed to digest the 3’-end region (within 16 bp of the stop codon) of the coding region of the target genes. The linearized sgRNA expression cassettes were then amplified by PCR from each gene’s modified pWS082 plasmid. Donor DNA cassettes were amplified from the plasmids in our previous screen,^11^ pAG25 (for DHFR-F[1,2] fragment) or pAG32 (for DHFR-F[3] fragment) using target gene-specific primers with 40 bp homology sequences as overhangs. The linearized Cas9 cassette was amplified from pWS176 by PCR. The three cassettes were gel purified by extracting fragments of the following sizes: sgRNA cassette, 1045 bp; Cas9 cassette, 9993 bp; donor DNA cassette for DHFR-F[1,2], 2009 bp, donor DNA cassette for DHFR-F[3], 1740 bp. The purified DNA cassettes were mixed in the following ratio and transformed: sgRNA cassette, 200 ng; Cas9 cassette, 100 ng; and donor DNA cassette, 5 μg. The average colony number was 21,879 per PPI (∼61-fold of the number of strain pooled barcodes (357)). The transformed cells were stored at −80°C for downstream processing. The number of strains in each pool (PPI) that received the mrDHFR fragments is shown in Figure 1F.

#### d) Adding a spike-in strains as reference for “no PPI”

As a reference for the growth rate of strains without mrDHFR-PCA complementation, two strains (#43 and #599, Table S2) were barcoded, but these were not tagged with the DHFR fragments. Each of these two strains were labeled by two unique barcodes to determine fitness replicability within each well. To perform the spike-in, the prepared library of tagged mrDHFR strains and the two reference strains were each incubated overnight at 30°C in YPD medium, and then adjusted to an OD_600_ of 2. We mixed 950 μL of the mrDHFR-tagged library pool and 12.5 μL of each reference strain. As a result, 5% spike-in of the reference samples were generated for all target PPI libraries, which were subsequently stored at −80°C for further processing. Altogether, the total number of unique barcodes in the pool is 361 (351 singly barcoded strains tagged with mrDHFR + (3 × 2) from 3 strains doubly barcoded tagged with mrDHFR + (2 × 2) from 2 strains doubly barcoded but untagged with mrDHFR) (Table S1).

### Selection and growth conditions for measuring PPI interactions

The constructed mrDHFR-tagged inbred strain libraries (61 PPIs + no-PPI reference) were grown as a serial batch culture in SD media (minimal synthetic defined bases, 2% glucose, 120 mg/L Leucine), 10 μg/mL methotrexate, and with (or without) drugs that served as “environmental” perturbations. The concentration of the drugs was determined in a prior GWAS study^26^ and PCA screen^12^: 5-FC (0.2 μM), fluconazole (100 μM), metformin (50 mM), and trifluoperazine (17.5 μM). Two biological replicates were passaged for all libraries and all drug conditions. For each sample, 40 μL of frozen cells (∼4 x 10^5^ cells) were inoculated into 760 μL media in 96 deep-well plates. The cells were grown at 30°C for 96 hours and passaged every 24 hours at a ratio of 1:8 (100 μL of cultured cells were added into 700 μL of fresh media). We measured the OD_600_ of the cultures at the end of each passage to calculate the cell number. After each passage, the deep-well plates were centrifuged at ∼1,900 × g for 10 minutes to pellet the cells. We stored cells from every passage at −20°C for the downstream analysis.

### Genomic barcode extraction and deep sequencing

The 61 PPIs measured in 5 environmental conditions (noDrug, 5-FC, fluconazole, metformin, and trifluoperazine), two timepoints (t = 0h and T = 96h), and two replicates resulted in 1,302 biological samples in twenty-one 96-well plates for downstream NGS analysis. To extract the gDNA, the cell pellets were resuspended in 300 μL of 50 mM EDTA containing 1 unit of Zymolyase (US Biological Life Sciences). We centrifuged the mixtures at 1,900 × g for 10 minutes after incubating them for 30 to 60 minutes at 37°C. By aspirating the supernatant, we removed the excess liquid. Next, we added 200 μL of lysis buffer (2% Triton X-100, 1% SDS, 100 mM NaCl, 10% Tris-HCl, 1 mM EDTA, pH 8), 2 μL of RNaseA, and 2 μL of protease K, and incubated for 10 minutes at 55 °C. The samples were vortexed, centrifuged at 1,900 x g for 10 minutes, and 200 μL of the supernatant was then transferred to fresh deep-well plates with 600 μL of 100% ethanol in each well. The samples were kept at room temperature for 15 minutes, then centrifuged at 1,900 x g for 10 minutes at 4°C, discarding the supernatant. We next obtained the isolated genome pellets, washed them using 150 liters of 70% ethanol, and centrifuged them at 1,900 x g for 10 minutes at 4°C. We removed the supernatant and let the pellet air dry at 60 °C for 5-10 minutes. The pellets were rehydrated for an hour at 65°C after being dissolved in 40 μL of TE. Strain barcodes were amplified from the extracted genomic DNA in a two-step PCR. The first PCR was performed using PrimeSTAR GXL polymerase (Takara) and 1:2000 SYBR Green I (20 μL/reaction) with 2 μL of extracted genomic DNA per reaction as a template. Because the concentrations of extracted genomic DNA varied, the number of PCR cycles was first determined by qPCR; the number of PCR cycles was chosen such that the amplification curve reached the mid to late log-linear phase (typically ∼24 cycles). The PCR amplification conditions were one cycle of 98°C for 3 min (initiation), followed by amplification cycles of 98°C for 10 sec, 55°C for 15 sec, and 68°C for 30 sec, repeated until the amplification curve reached the mid to late log-linear phase. Primers used for this reaction are of the following format:

Forward: TCGTCGGCAGCGTCAGATGTGTATAAGAGACAG-*NNNNNNNN*-AGGCGGTTTGCGTATTGGGC

Reverse: GTCTCGTGGGCTCGGAGATGTGTATAAGAGACAG-*NNNNNNNN-* ACGGCACGCATCTTTGGCTG

The *N*s corresponds to in-house sample indices, allowing the unique labeling of samples in each well of the 96-well plate during multiplexed Illumina NGS.

The PCR products were pooled per growth media condition and purified with DNA Clean & Concentrator (Zymo Research). The second PCR was performed using PrimeSTAR GXL polymerase (Takara) with 5 ng of purified product from the previous PCR was used as a template (50 μL/reaction). Primers were unique pairs of index primers (i5 and i7) from the Nextera XT DNA library preparation kit (Illumina). The PCR amplification conditions were one cycle of 98°C for 45 sec, then 12 cycles each of 98°C for 10 s, 55°C for 15 s, and 68°C for 30 s. The second PCR products were purified using Agencourt AMPure XP PCR purification kit (Beckman Coulter). We performed pair-end sequencing on Illumina NextSeq platforms with 30% balanced DNA (phiX).

### Reverse transcriptase real-time quantitative PCR (RT-qPCR)

The selected strains from wine or clinical backgrounds were grown in synthetic complete medium at 30°C, overnight. Cells were collected by centrifugation and total RNA was extracted using TRIzol reagent (Invitrogen) following manufacturer’s instructions. To prepare cDNA, approximately 2 µg of total RNA was reversed transcribed using All-In-One 5X RT MasterMix (abm). The reaction products were diluted 10 times before using them for amplification. qPCR reactions were performed using FastStart Essential DNA Green Master (Roche Diagnostics) in a LightCycler 96 Real-Time PCR System (Roche Applied Science). All reactions were performed in triplicates. The amplification program was set at an initial step of 6 min at 95°C, 50 amplification cycles of 20 sec at 95°C, 20 sec at 58°C and 20 sec at 72°C, followed by a high resolution melting from 60°C to 98°C. Primers for *EGR11* (ERG11-F1: GATTCCATGGGTCGGTAGTGC, ERG11-R1: CACAGTCATGACTCTTCCTAAC) and *SUT153* (SUT153-F1: CAGGCATAAAATCGATGTCTCCT, SUT153-R1: CTGGCCCTAAACGAAACGAAAC) target genes were designed in this study while primers for the reference *ACT1* gene were used before in Brickner *et al*.^91^ Gene expression levels for the two loci relative to the control gene were determined with LightCycler 96 (software version 1.1).

## QUANTIFICATION AND STATICAL ANALYSIS

### FASTQ demultiplexing and pre-processing

Our NGS run resulted in >350M reads with 89% at Phred score >= 30. Using *AdapterRemoval* (version 2.3.2),^92^ we allowed at most 2 mismatches within the inner-adapters (--barcode-mm-r1 = 2 and --barcode-mm-r2 = 2) to account for mutations that could arise from PCR and/or NGS. Since the in-house sample indices have Hamming distances greater than 2, ∼98% of the reads uniquely mapped to their biological samples.

### Barcode extraction and mapping to inbred strains

We extracted the strains barcodes from the reads using the command *bartender_extractor_com* of *Bartender* (version 1.1)^93^ by providing the following regular expression: *TGGGC[5]CAGGTCTGAAGCTGTCGCAC[5]GAAAT*, where TGGGC (preceding sequence) and GAAAT (succeeding sequence) that frame the construct double barcodes. The two 5-nucleotide barcode regions are linked by a constant spacer (CAGGTCTGAAGCTGTCGCAC). Additionally, we allowed 2 mismatches within the preceeding and succeeding sequences to account for mutations during PCR or/and sequencing steps. This step resulted in 10-nucleotide barcode reads, which we mapped to the 361 unique strain barcodes (Table S2) using in-house Python scripts. Since the Hamming distance between our strain barcodes is >=3, we allowed for at most 1 mismatch between the barcode and strain IDs and successfully mapped 98% of the barcodes.

### Fitness and PPI estimation

In each well containing the pooled strains but tagged for a specific PPI π (Figure 1C(iii)), we estimated the relative fitness of a strain *i* as the log2-fold-change (LFC) of its normalized frequency *n* between initial and final timepoints (0h and 96h, respectively):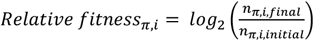. We compared the fitness estimate between two replicates for all strains across all conditions (∼223,820 fitness values) using a density plot in *Matplotlib* Python library (Fig. 1F). The density values were estimated using Gaussian kernels. To estimate the PPI strength for each bait/prey π in strain *i*, we subtracted the average fitness of the reference strains (table S2) that were untagged with DHFR fragments:

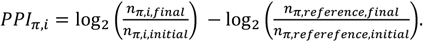

### piQTL association mapping

#### (a) Encoding of the genotype matrix and calculation of residual linkage disequilibrium

The inbred strains were previously genotyped by She and Jarosz.^26^ They provide the genotype matrix containing 12,054 SNPs, where 1 denotes the wine parental haplotype, −1 the clinical parental haplotype, and 0 is unphased haplotype which we consider as missing genotype. Using *snpStats*, we calculated amount of linkage disequilibrium (LD) between all pairs of SNPs as the Pearson correlation between their genotype values across all 357 strains. LD blocks were defined as genetic regions with neighboring SNPs with LD > 0.75.

#### b) piQTL analyses

Our first piQTL approach (approach 1) relied on a simple linear regression model implemented in *Matrix eQTL*^35^, where instead of providing a gene expression matrix as inputs for the phenotyping, we provided our PPI quantification matrix. The association between the changes of PPI and genotype was assumed to be linear: 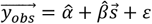, where 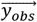 is the vector of measured PPI values across all strains,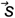 is vector of strain genotype at a SNP locus, 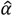 is the intercept, 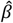 is the slope (QTL “effect size”) and *ε* is a random variable from independent and normal distribution. This was followed by calculation of a *t*-statistic test that provides a *P*-value. Due to the high number of association tests performed (61 PPIs × 5 environments × 12K SNPs), a false discovery rate (FDR) was computed to account for false positive associations.

Our second piQTL approach (approach 2) modeled the relationship between the genotype and the a phenotype (PPI) using a generalized linear model implemented in *rMVP*^94^ as follows: 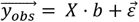, where 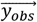 is a vector of observed PPI values across all strains, *X* is a matrix of fixed effects and testing SNPs, *b* is an incidence matrix for *X*, and 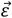 is a vector of residual that follows a normal distribution with mean of zero. A crucial distinction between the first approach is that we incorporated as a covariate in *X*, the kinship matrix inferred from the genotypes. This means that the hypothesis testing accounts for residual LD that might exist between the SNPs.

We performed both approaches on all SNPs (12,054) and for all PPI under different environmental conditions. SNPs that were known to be QTLs for methotrexate in the previous GWAS study^26^ were masked in the analysis. SNPs are considered piQTLs if they were significant (FDR < 0.05) in *both* approaches.

#### c) Heatmap of colocalized piQTLs across 61 PPIs

For each environmental condition (no Drug, 5-FC, fluconazole, metformin, trifluoperazine), we built a matrix 62 PPI rows x 12,054 SNP columns where we assigned a value of 1 for SNPs defined as a piQTL under the presence of the drug (FDR < 0.05 which corresponds to *P*-value < 0.0001) or its FDR changed by an order of magnitude between the presence and absence of the drug (Figs. 3A and 5A). For the visualization, SNPs positions with no piQTL across all the PPIs are not shown. The 61 PPIs were ordered based on their proximity in the PPI network defined from a genome-wide *in vivo* yeast protein interactome.^11^ Proximity on the network was calculated as the shortest weighted path between nodes, where the weight is the *in vivo* PPI strength.^11^ This calculation is implemented in *NetworkX*.

#### d) Interactive visualization of piQTLs

We provided two *Shiny/R* applications to dynamically visualize the results of the piQTL mapping. Both webtools can be run locally after installing the required library from R (version 4.2.2). These web interfaces allow to visualize the piQTL graphs, such as Manhattan, quantile-quantile and volcano plots, and to browse the piQTL results across the yeast annotated genome and several genome annotations. They are both available on the piQTL project webpage (https://ladyson1806.github.io/SerohijosLab-piQTL/).

##### Manhattan, quantile-quantile (QQ) and volcano plots

Before visualization, the MTX-specific piQTLs were excluded. MTX-specific piQTLs were defined as piQTLs that appear with MTX treatment, both in the presence and absence of the four environmental drug conditions. These piQTLs corresponded to 45 SNPs, all located in chromosome 15.^26^ The Manhattan plot display all *P*-values calculated from the association’s tests using the generalized linear model (approach 2). The quantile-quantile (Q-Q) plot represents the distribution of theoretical versus experimental *P*-values. These graphs were generated by using *Manhattanly*, an R library (https://github.com/sahirbhatnagar/manhattanly/, version 0.3.0).

##### Interactive genome browser

This *R/Shiny* app is a custom wrapper to load the piQTL Manhattan plots on the annotated yeast genome (version *sacCer3* from the Broad Institute). Clicking on the hits will yield information on the annotations of a SNP (SNP ID, chromosome ID, base-pair position, information of their SNP class, locus ID, gene/genome feature ID, and SGD ID) and piQTL association statistics calculated from the rMVP analyses (*P*-value, effect size, standard error). Additional information is plotted, such as locations of non-coding RNAs^38,39^ CUT, SUT and XUT, as well as residual LD blocks (LD score > 0.75). These graphs were generated by using the R library *igvShiny* (https://github.com/paul-shannon/igvShiny, version 1.5.90009)

### Mapping between human and yeast homologs

To correlate the yeast piQTL hits of the human drugs metformin and trifluoperazine, we assigned the yeast genes to their human homologs using the orthology mapping from Alliance of Genome Resources (https://www.alliancegenome.org/downloads, Alliance Combined Orthology Data), which combines predicted and manually curated annotations. We used the “Stringent” option.

### Mapping piQTLs biological networks

PiQTLs of a given PPI were projected onto the yeast proteome-wide *in vivo* PPI network by Tarassov *et al*.^11^ and onto the genome-wide genetic interaction network by Costanzo *et al*.^45^. Shortest distance (smallest number of edges) between piQTLs on the network were calculated in *NetworkX*. To test if the distances between piQTLs of a PPI are significantly shorter (or longer) than random, we performed 10,000 simulations where *n* protein nodes (with SNPs) were randomly picked and then calculated the shortest distances between them. *n* is equal to the number of piQTLs for a PPI. Kolmogorov-Smirnoff (KS) test was performed to compare the distribution of shortest distances of piQTLs and the random distribution.

### Meta-analysis of piQTLs from 61 PPIs

We performed meta-analysis using *Plink*^95^ (https://zzz.bwh.harvard.edu/plink/) starting from the summary statistics generated by our piQTL analysis of the 61 PPIs. The simplest model for meta-analysis is the “fixed-effects model” that assumes that the independent association studies are measuring the same phenotypic effect of the SNP at locus *l*, which can be estimated by 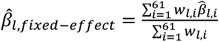, where 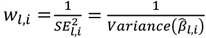 is the inverse variance (“precision”) of the effect size 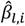 for protein *i* at SNP *l*. Although, the cloning of the mrDHFR fragments and screens are independent experiments, the 61 PPIs are not identical, and hence, not expected to have the same (or “fixed”) effect size. This assumption is relaxed in the random-effects model of meta-analysis, where instead the values 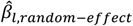 are assumed to follow a normal distribution. Results from both fixed- and random-effects models are provided, but we report the *P*-values arising from the random-effects model (Tables S3, S4, S5, and S6).

## DATA AND SOFTWARE AVAILABILITY

All computational analyses are available in *GitHub* (https://github.com/ladyson1806/Serohijoslab-piQTL). Raw sequence reads are available at https://www.ncbi.nlm.nih.gov/geo/ (GSE246414).

## SUPPLEMENTAL INFORMATION

**Figure S1.**
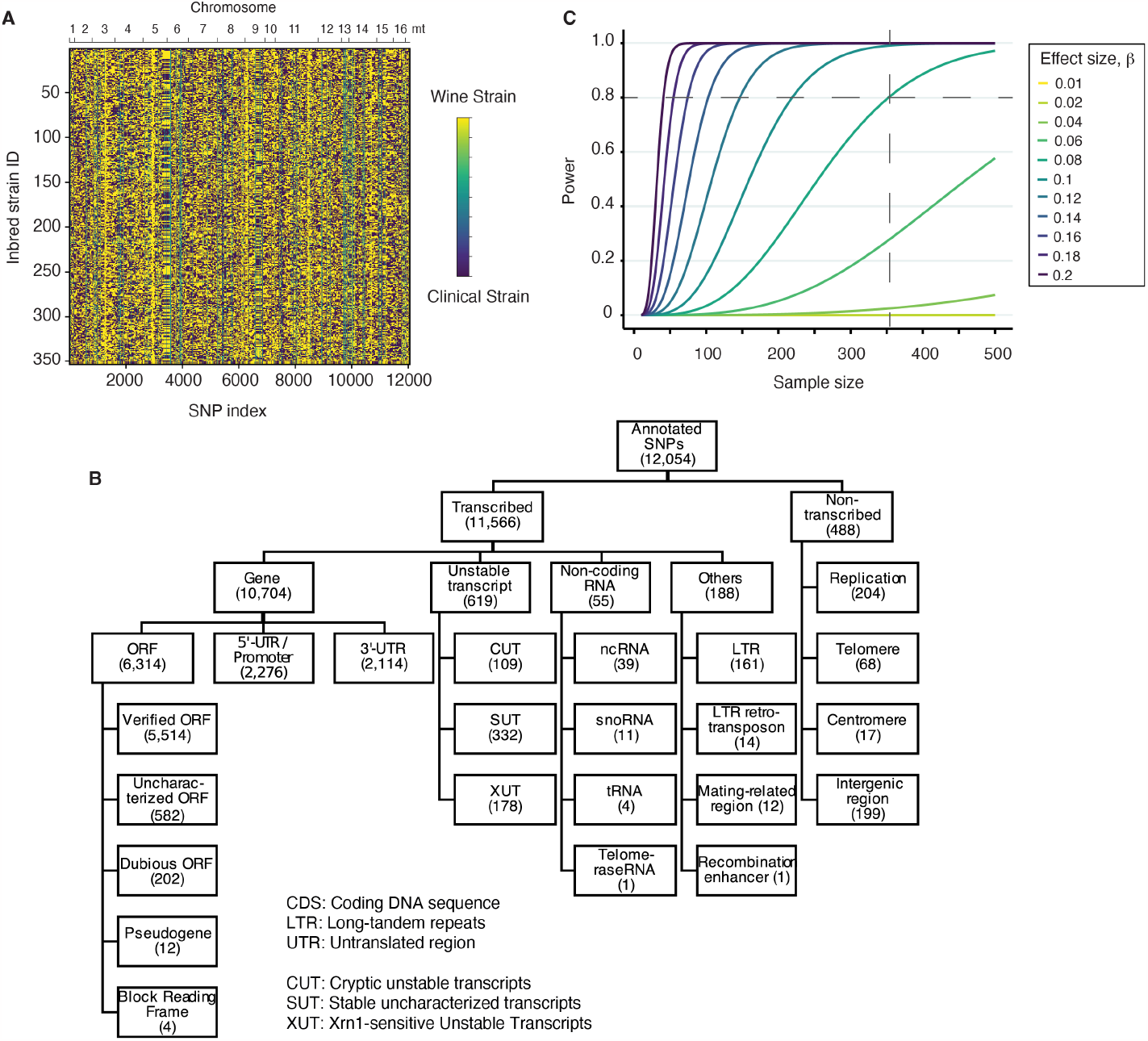
Genotype map of inbred strains and genomic architecture of ∼12K mutations, related to Figure 1. **(A)** Genotyping of the 353 inbred strains from a previous study by She & Jarosz^26^ shows ∼12K SNPs across all yeast chromosomes and mitochondrial DNA. **(B)** Distribution of the ∼12K SNPs across the functional and architectural features of the yeast genome. Recent annotation of *S. cerevisiae* S288C was used as a reference.^88^ **(C)** Power calculation estimate showing that a sample size of 354 strains can detect effect sizes ∼0.06 or higher (estimation used minor allele frequency = 0.5 since mutations segregate at 50:50 because of inbreeding).

**Figure S2.**
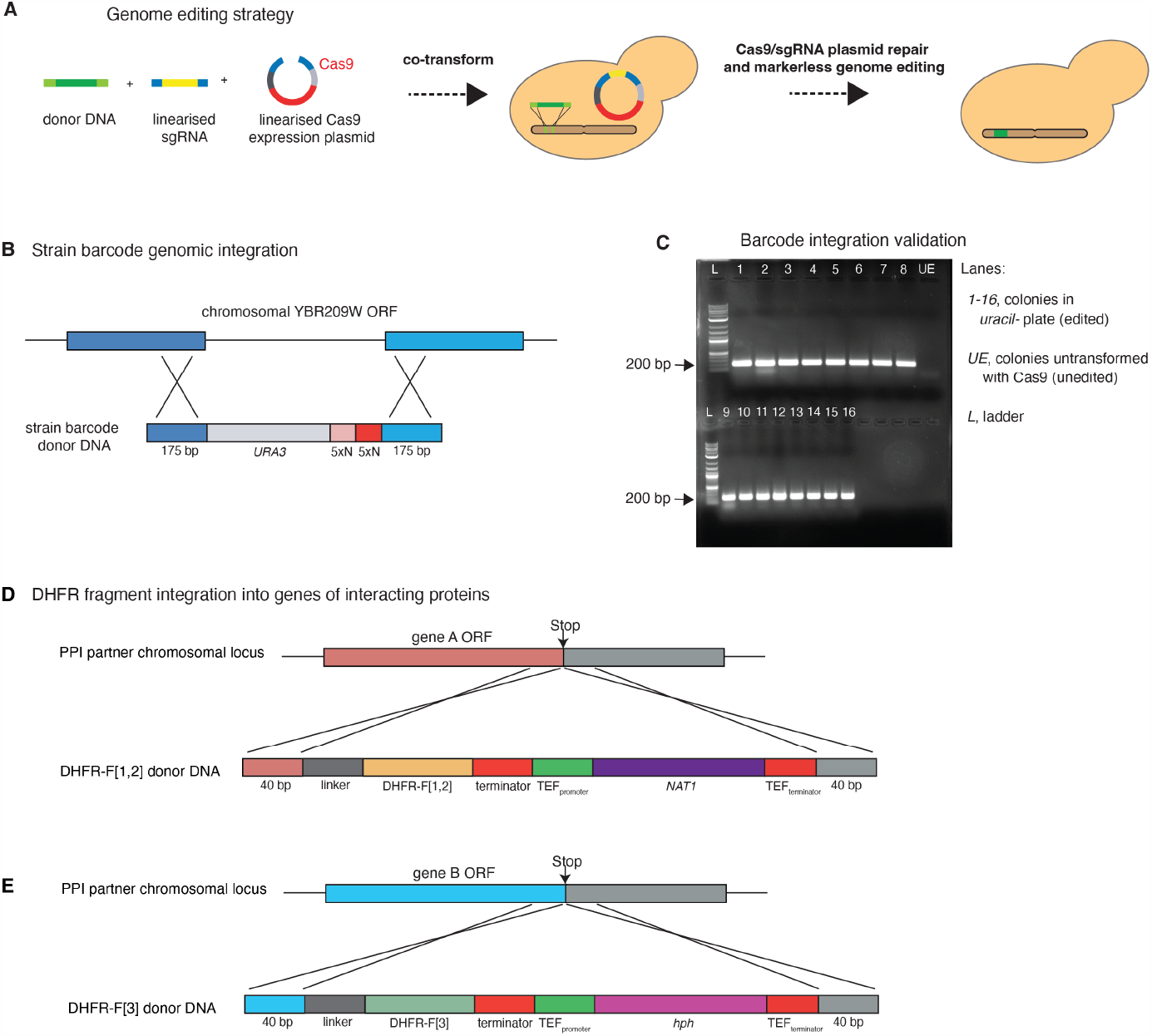
Genome-editing strategy for strain barcoding and tagging gene pairs with DHFR fragments, related to Figure 1. **(A)** High-efficiency yeast genome editing that couples CRISPR/Cas9 with homologous recombination.^89^ A “donor DNA” with homology arms to the target genomic locus, an sgRNA targeting the locus of integration, and a linearized Cas9 expression vector are co-transformed into yeast. Gap repair leads to the reconstitution of the Cas9 expression plasmid, followed by the cutting of the genomic target locus by Cas9/sgRNA, and subsequent repair by homologous recombination. **(B)** To perform the chromosomal barcoding, the donor DNA contains two NNNNN sites (denoted as 5×N in the diagram) that uniquely label each of the 354 strains (table S2). This donor cassette is in tandem with a *URA3* gene selection marker that allows selection on minimal media without uracil. The sgRNA targets for barcode integration of a neutral locus in the genome (YBR209W).^34^ **(C)** We validated the barcode integration by growing the cells in the absence of uracil. Only cells that successfully integrated the *URA3* gene survived. We picked 16 random colonies and performed colony PCR to amplify a 215 bp-region spanning the insert and the integration locus, which showed the expected product size. **(D** and **E)** The genome editing for DHFR fragment tagging follows panel A. Panel D is the donor cassette for DHFR-F[1,2] consisting of a 40 bp homologous sequence on the left, DHFR-F[1,2], terminator, followed by TEF promoter, nourseothricin N-acetyl transferase (*NAT1*) which confers resistance to nourseothricin, TEF terminator and finally a 40 bp homologous sequence on the right. The linker encodes a 10 amino acid (Gly-Gly-Gly-Gly-Ser)_2x_ connection between the protein being tagged and the DHFR reporter fragment. The donor cassette for DHFR-F[3] (panel E) consists of the same elements, except for a different selection marker, hygromycin B phosphotransferase (*HPH*), which confers resistance to hygromycin B. The sequence of the sgRNA and homology arms for each gene are listed in Table S1.

**Figure S3.**
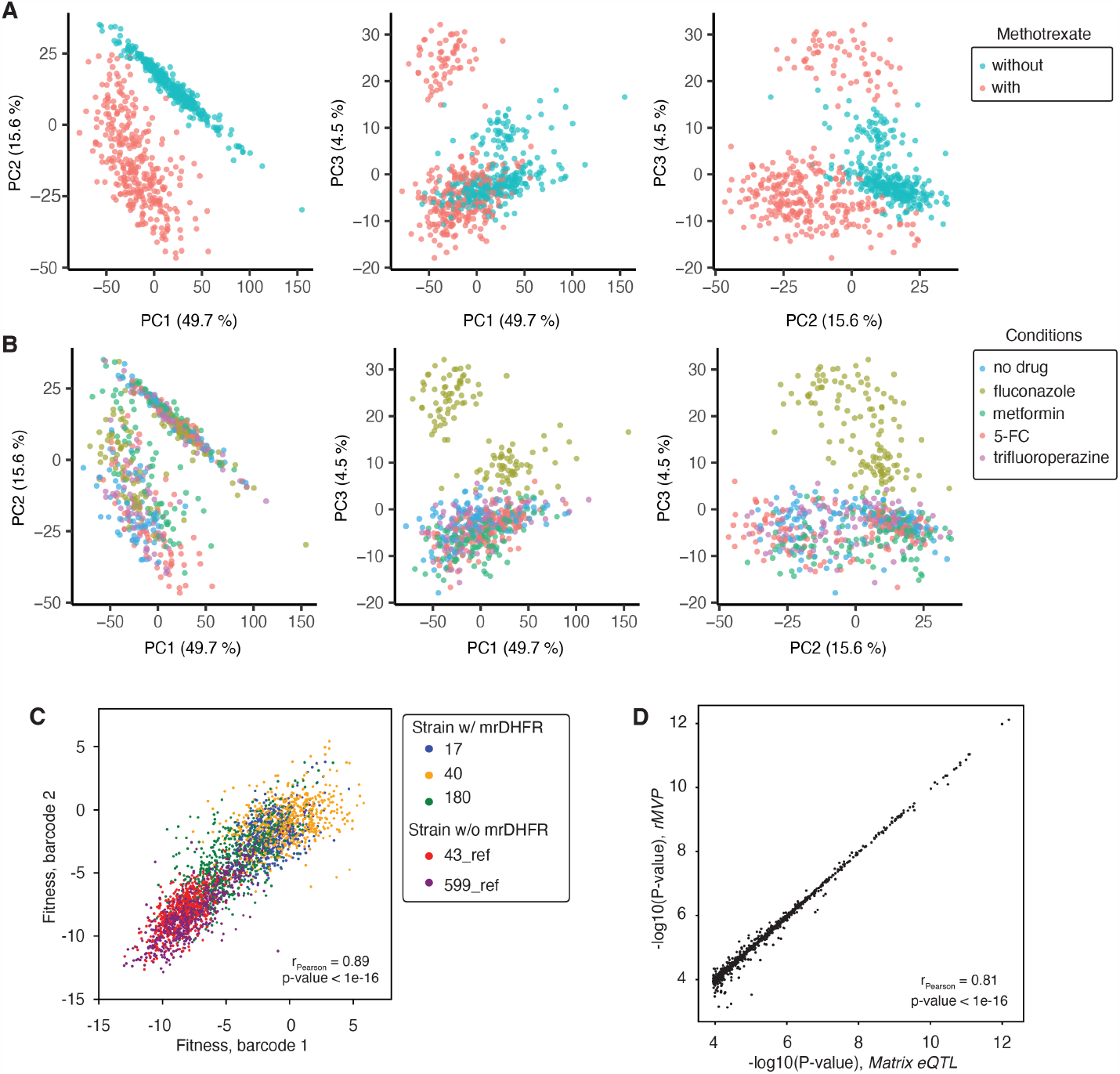
Replicability and principal component analyses for the PPIs, related to Figure 1. **(A** and **B)** Principal component analyses for the PPIs measured across different conditions. Projection of the 62 PPI (each point as an average over 354 strains) onto the first three principal components. Colors correspond to with and without methotrexate treatment (panel A). Panel B is similar to panel A, but the colors correspond to different drug conditions. **(B)** Within-well replicability of fitness estimates from doubly barcoded strains. Three inbred strains (#17, #40, and #180) were independently labeled with two barcodes prior to being pooled and tagged with the mDHFR fragments. Two other strains (#43 and #599) were also labeled with two barcodes but were not tagged with mDHFR. The fitness of the latter strains served as reference for “no PPI” or fitness in the absence of mrDHFR complementation. Shown is the correlation of the fitness estimates from the two unique barcodes under methotrexate and drugs conditions of the 61 PPIs. Expectedly, strains without mrDHFR complementation have the lowest fitness under MTX treatment. **(D)** We perform piQTL mapping using *matrix eQTL*^35^ without correction for linkage disequilibrium and *rMVP*^94^ that corrects for LD using the correlation in the genotypic matrix. Due to the minimal presence of LD in the inbred population cohort, the association mapping with and without LD correction are strongly correlated. We only consider as piQTL those SNPs that have FDR < 0.05 (P-value∼10^-3.9^) in *both* approaches.

**Figure S4.**
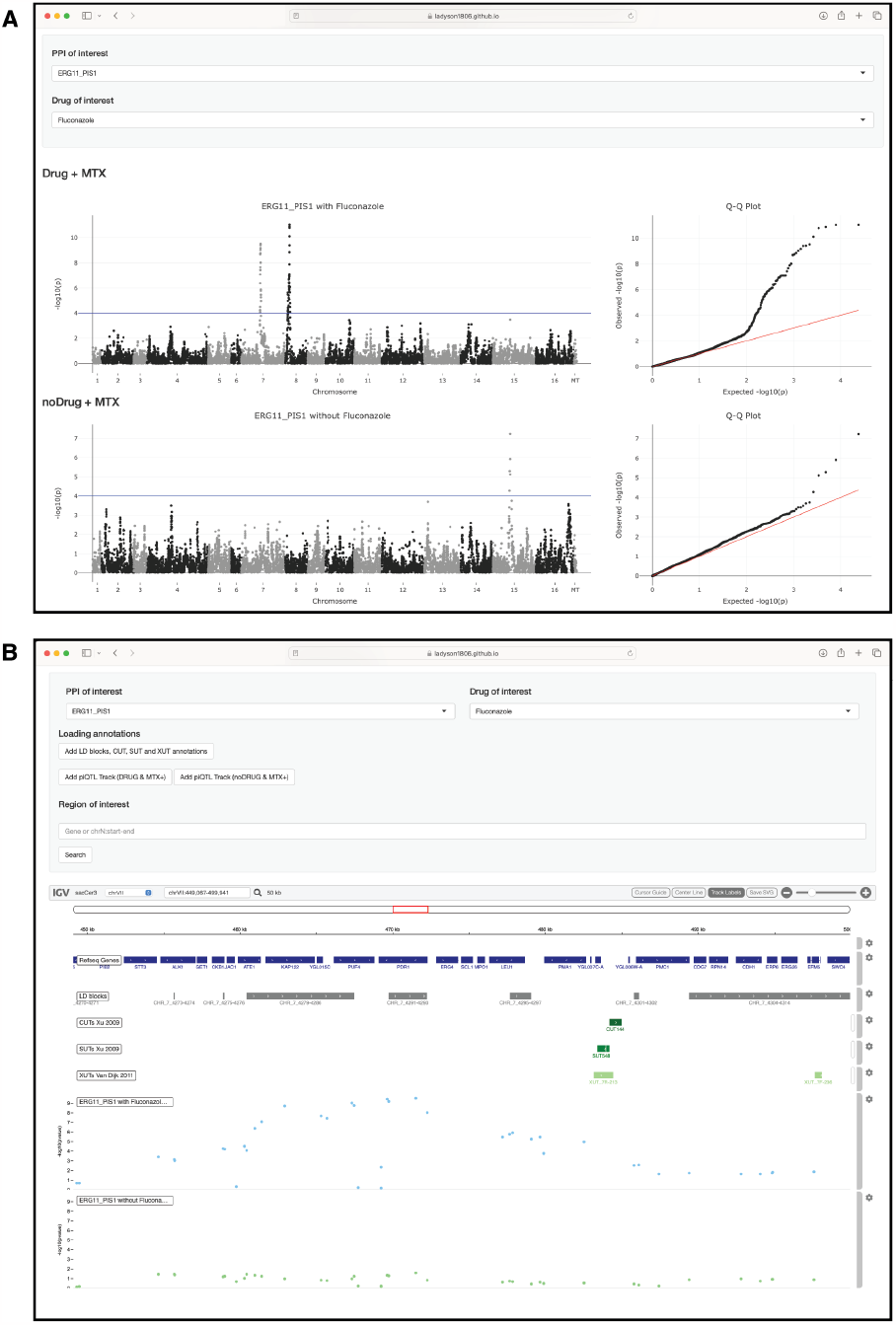
Interactive piQTL webserver, related to Figures 2 and 3. **(A** and **B)**. All PPI genome-wide association analysis results are available in an interactive server (https://ladyson1806.github.io/SerohijosLab-piQTL/) showing the statistical significance of the association and Q-Q plots (panel A). A genome-browser (panel B) also provide details of the genomic features; annotations of non-coding RNAs SUT, XUT, and CUT; and residual LD blocks (LD r^2^> 0.75).

**Fig. S5.**
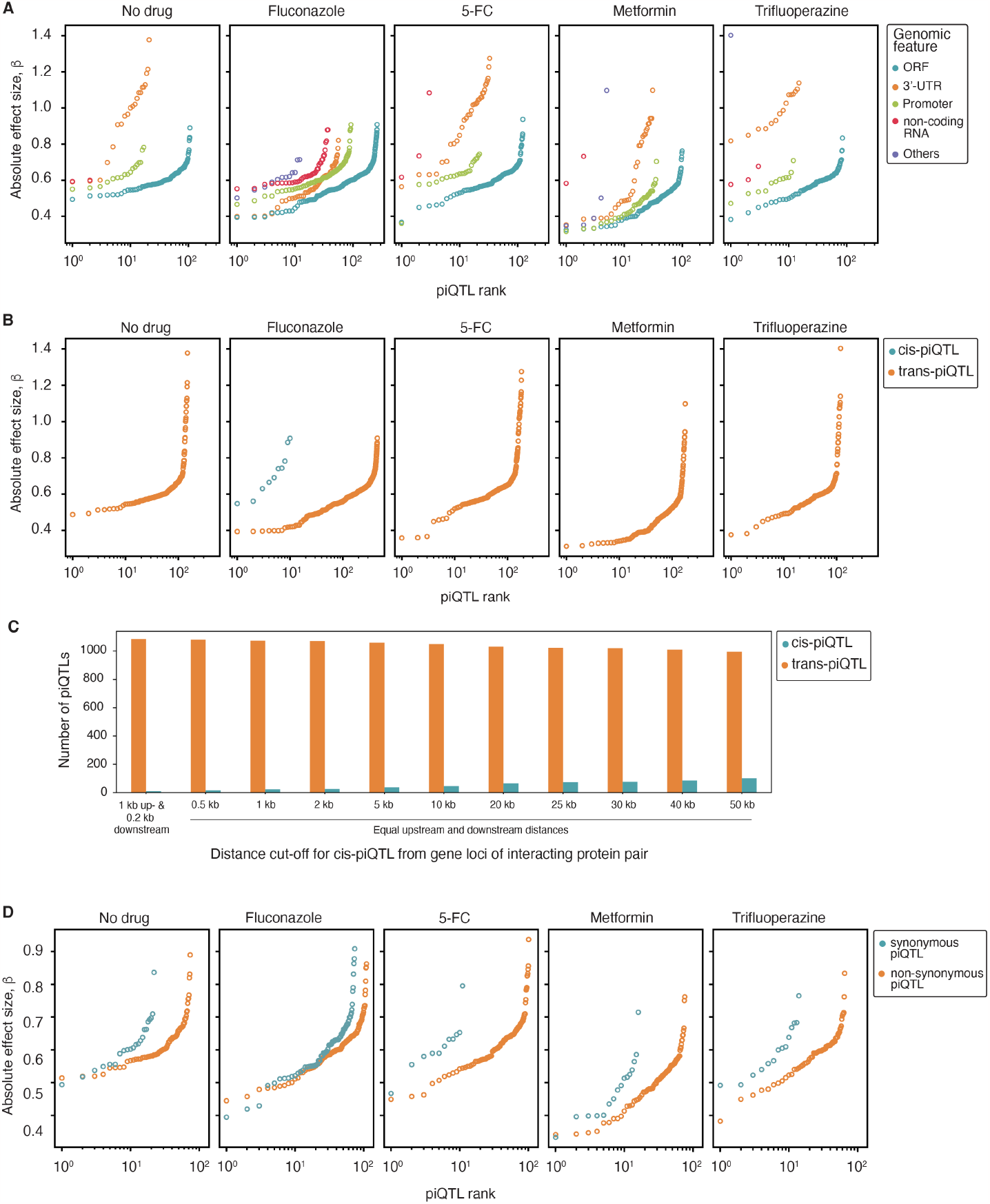
piQTLs across genomic and sequence features, related to Figure 3. **(A)** Effect sizes of largest piQTLs accounting for existence of residual LD blocks not broken by inbreeding. Similar to Figure 2A, but only the most significant piQTL in an LD block is considered. LD block is defined as regions with SNPs that have contiguous LD r^2^ ≥ 0.75, as previously done.^36^ Counting only the most significant piQTL in an LD block yield 1,096 piQTLs in 290 unique SNP loci. **(B)** Cis- and trans-piQTLs across drug conditions. Only the most significant piQTL in an LD block is considered. Note that cis-piQTLs are detected only under fluconazole. **(C)** Number of cis- and trans-piQTL as a function of distance from the gene loci of the protein pair forming the PPI. The first distance criteria for cis-piQTL (within 1 kb upstream or 200 bp downstream) follows from a previous yeast genome-wide eQTL study.^40^ Trans-piQTL significantly outnumber cis-piQTL even when the cut-off distance for cis-piQTL is increased. **(D)** Synonymous and non-synonymous piQTLs in protein-coding regions. The effect sizes between the two groups are comparable across all conditions (KS-test p-values testing for difference in distribution of effect sizes were insignificant). Only the most significant piQTL in an LD block is considered.

**Figure S6.**
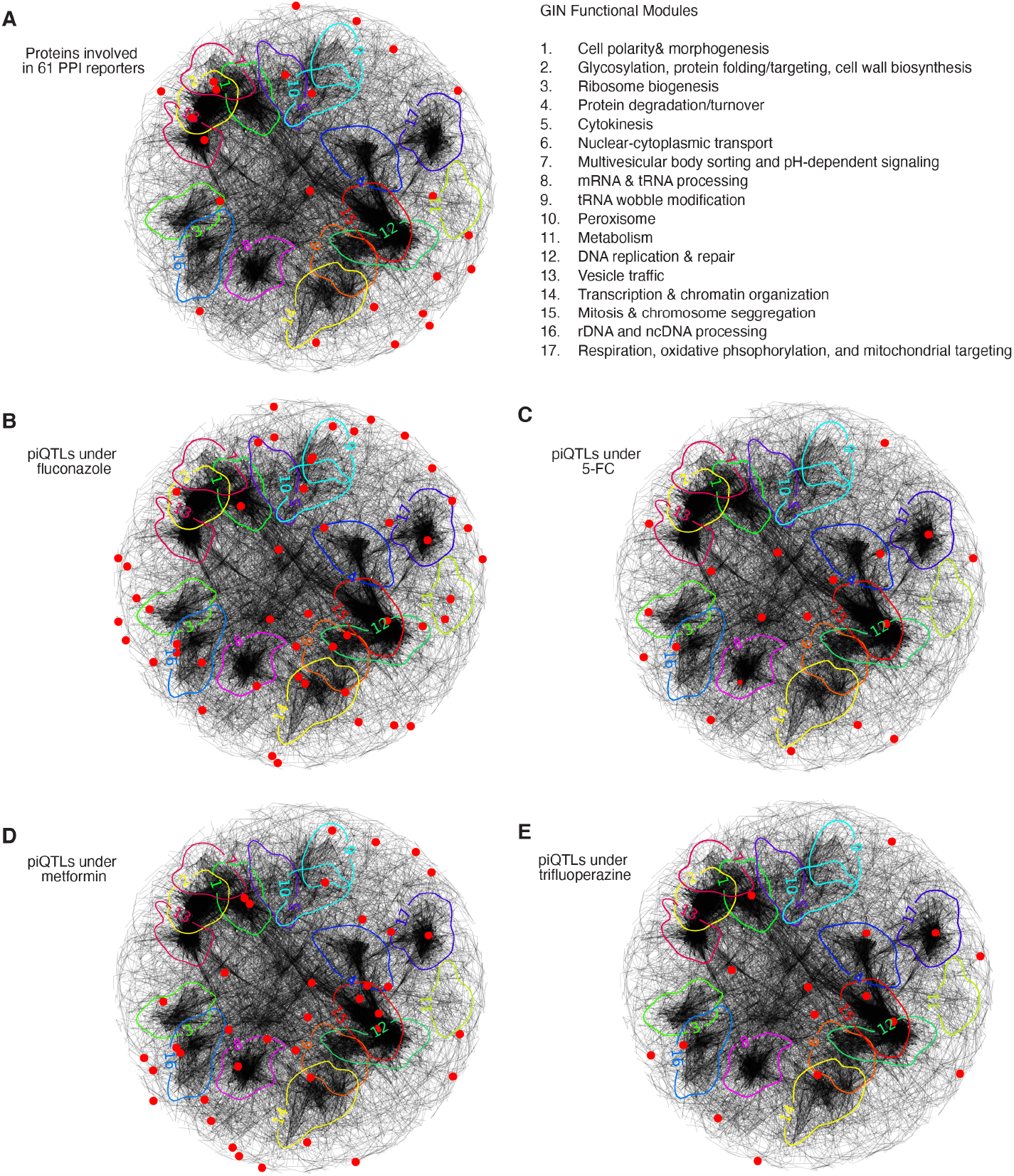
piQTLs on the genetic interaction network (GIN) defined by double-gene knockouts, related to Figure 4. **(A)** GIN network location of 44 proteins (red dots) participating in the 61 reporter PPIs. Functional modules as defined by Costanzo et al.^45^ are highlighted. **(B** to **E)** piQTLs (red dots) on the GIN for the different drug conditions.

**Figure S6.**
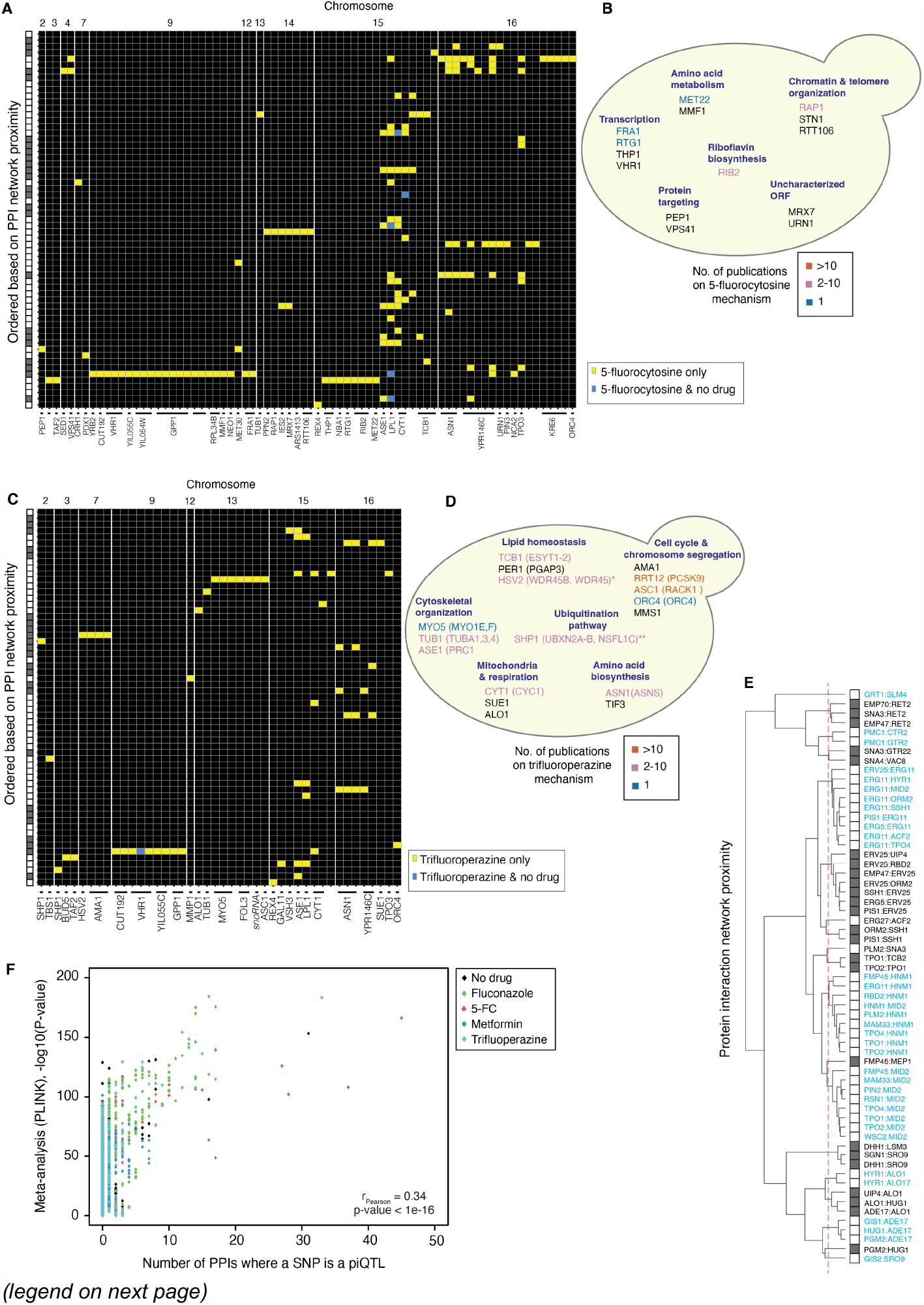
piQTLs for 5-FC (5-fluorocytosine) and piQTLs for trifluoperazine, related to Figure 4 and 7. **(A)** SNPs that were piQTLs under 5-FC for at least one PPI are shown in panel A. Yellow boxes are piQTLs identified only upon 5-FC treatment. Blue are piQTLs in both with and without 5-FC, but whose FDR changed by an order of magnitude. Rows are ordered according to panel E. Gene descriptions are in Table S4. **(B)** Biological functions of piQTLs under 5-FC. Gene name colors correspond to number of studies in PubMed relating the same gene to mechanisms of 5-FC and antifungal resistance. **(C)** Similar to panel A but for piQTLs under trifluoperazine. Gene descriptions are in Table S6. **(D)** Biological processes of trifluoperazine piQTLs, where the human homolog(s) of the yeast proteins are shown in the parenthesis. **(E)** Hierarchical clustering of the 61 PPIs based on the number of edges separating any two pair in the PPI network (Figure 2B). The hierarchical tree and the PPI groups (rightmost bar) define the y-axes of the heatmaps in Figures 4 and 7. **(F)** Meta-piQTL analysis. Significance from meta-analysis using a random-effect model (implemented in *PLINK*^95^) is higher for SNPs that are piQTLs in multiple PPIs.

## Notes

https://github.com/ladyson1806/Serohijoslab-piQTL

### Competing Interest Statement

The authors have declared no competing interest.

